# A novel data filtering method resolves the controversy in the phylogeny of the Chondrichthyes

**DOI:** 10.1101/2025.08.19.671162

**Authors:** Junman Huang, Michael Hofreiter, Leslie Robert Noble, Nicolas Straube, Gavin J. P. Naylor, Chenhong Li

## Abstract

Phylogenomics, which uses genome-scale data for phylogenetic inference, has clarified many controversial nodes in the tree of life. Such extensive data improve tree resolution and better reflects organismal history compared to analyses based on single or a few genetic loci. However, some relationships within the tree of life remain unresolved, as increased data can yield high node support without ensuring accuracy due to systematic errors. For example, the order-level relationships among chondrichthyans are still contentious despite the use of phylogenomic data. To address systematic errors, complex models have been developed, and filtering for less erroneous data shows great promise. Current metric-based filtering methods rank loci based on overall tree statistics, but problematic signals are often local; and topology-based data filtering approaches struggle with circular assumptions. In this study, we introduced two novel metric-based data filtering methods based on the ratio of local branch length or GC content between problematic clades. We applied these methods to a dataset of 4,452 single-copy exons extracted from 98 chondrichthyan species. The results using all loci showed that the Hexanchiformes was positioned at the root of Elasmobranchii, pulling other squalomorphs to the basal position and rendering Squalomorphii paraphyletic. Contrastingly, filtering for loci with more even branch length, the branch ratio method (absRatioLen) strongly supported the monophyly of all superorders of the chondrichthyans as well as their higher classification grouping, such as Selachii and Batoidea. By concentrating on problematic nodes, our assumption-free filtering methods demonstrate significant potential in resolving contentious relationships in the tree of life.

## Introduction

The advent of genome-scale data together with innovative statistical methods has revolutionized phylogenetic inference, resolving numerous previously contentious branches of the tree of life (Steenwyk et al., 2023). Despite these advances, persistent challenges arise from systematic errors, such as base composition bias, substitution rate heterogeneity, and ancient hybridization, confounding the resolution of rapid evolutionary radiations (Schrempf et al., 2020; Stiller et al., 2024). Even with expansive genomic datasets, such errors often obscure deep phylogenetic relationships, necessitating strategies to enhance signal-to-noise ratios (Jeffroy et al., 2006; Philippe et al., 2011). Central to this effort is the filtration of datasets to retain loci with robust phylogenetic signal while excluding those prone to homoplasy or compositional bias (Arcila et al., 2017; Li et al., 2012; Shen et al., 2017).

Filtration strategies fall into two categories: metric-based filtering, which selects loci based on inherent properties such as evolutionary rate, base composition uniformity, or molecular clock-likeness (Kuang et al., 2018; Li, et al., 2012; Steenwyk et al., 2021); and topology-based filtering, which evaluates loci against alternative phylogenetic hypotheses, such as gene genealogy interrogation (GGI) (Arcila et al., 2017), or gene-wise likelihood scores (GLS) (Shen et al., 2017). For metric-based filtering, traditional metrics like relative composition frequency variability (RCFV), calculated at the gene level, often mask taxon-specific variation (Zhong et al., 2011). That is, localized systematic errors within specific clades can be obscured when measured across broad taxonomic scales. For example, Li et al. (2012) found that using conventional metrics (*p*-distance and RCV) to filter genes was insufficient to address systematic biases in resolving the phylogeny of chondrichthyans. Conversely, topology-based filtering frameworks remain vulnerable to subjectivity, particularly when relying on maximum likelihood (ML) trees as reference topologies, a practice that risks circular reasoning. These drawbacks underscore the need for objective, data-driven approaches focusing on problematic nodes to evaluate the phylogenetic signal of each locus.

Chondrichthyes (sharks, rays, skates, and chimaeras) originated in the Paleozoic era (∼460– 300 million years ago) and have diversified into various ecological niches and forms (Heinicke et al., 2009). Extant chondrichthyans are broadly classified into two major groups: Holocephali (chimaeras) and Elasmobranchii (sharks, rays and skates), which is further divided into two clades: Selachii (sharks), comprising Galeomorphii and Squalomorphii, and Batoidea (rays and skates) (Nelson et al. 2016). Although the taxonomic classification of chondrichthyans in general is established, phylogenetic relationships among major lineages, particularly Galeomorphii, Squalomorphii, and Batoidea, remain controversial, and interrelationships within Batoidea are also under continued debate (Fig. 1).

**Fig. 1.**
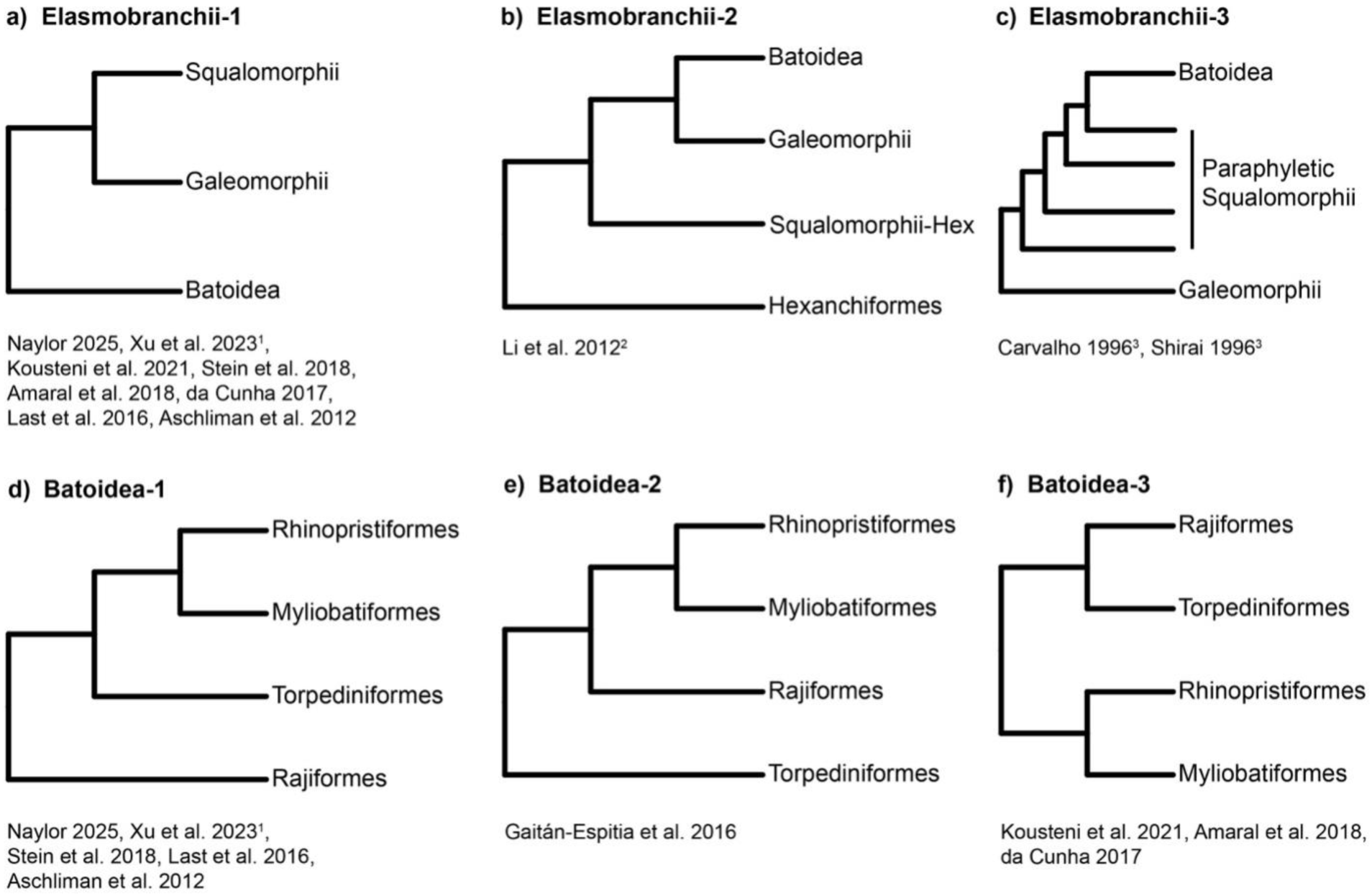
Summary of phylogenetic hypotheses for elasmobranch orders from 12 studies: (a-c) three alternative topologies within Elasmobranch, (d-f) three proposed topologies within Batoidea. ^1^phylogenies inferred using genomic exons. ^2^topology recovered based on nuclear datasets. ^3^studies based exclusively on morphological data. The remaining studies are based primarily on mitochondrial loci.

Much of the molecular phylogenetic work on Chondrichthyes has relied heavily on mitochondrial genes or a limited number of nuclear markers (e.g., RAG1) (Amaral et al. 2018; Cunha et al. 2017; Kousteni et al. 2021; Last et al. 2016; Naylor, 2025; Aschliman et al. 2012; Stein et al. 2018). These studies typically support the monophyly of both Galeomorphii and Squalomorphii (Fig. 1a), but mitochondrial DNA, being maternally inherited, represents only a single genealogical history and may not reflect the species tree. Intriguingly, a study using ten nuclear genes recovered an unprecedented topology in which Galeomorphii clustered with Batoidea rather than Squalomorphii (Li et al., 2012; Fig. 1b), a result not easily reconciled with morphological data. A more recent phylogenomic study including thousands of loci recovered a conventional topology in which Selachii is monophyletic (Xu et al., 2023), but notably excluded Hexanchiformes, a key lineage found to disrupt ingroup–outgroup resolution in Li et al. (2012). In addition to molecular hypotheses, a classic morphological hypothesis, the Hypnosqualean model, proposes Squalomorphii as paraphyletic and as sister to Batoidea (Carvalho, 1996; Shirai, 1996; Fig. 1c).

Relationships within Batoidea are also contentious. Three competing topologies have emerged from different data sources (Fig. 1d–f): most nuclear and mitochondrial studies support Rajiformes as the basal batoid group with the Torpediniformes sister to Rhinopristiformes + Myliobatiformes (e.g., Naylor, 2025; Xu et al., 2023; Stein et al., 2018; Aschliman et al., 2012), while mitogenomic datasets often support alternative relationships (Amaral et al., 2018; Cunha et al., 2017; Gaitán-Espitia et al., 2016; Kousteni et al., 2021; Fig. 1e-f). Morphological evidence further complicates this picture, with some studies placing Rhinopristiformes as the most basal batoid lineage (Villalobos-Segura et al., 2022).

Given unresolved conflicts in chondrichthyan evolution, we developed a phylogenomic framework optimized for large genomic datasets, using a matrix of 4,452 single-copy exons generated through target capture and genome mining across 98 chondrichthyan species. To improve phylogenetic signal, we introduce two novel filtering strategies that quantify locus-specific biases associated with problematic nodes. The first, branch length ratio filtering (absRatioLen), selects loci with balanced evolutionary rates across focal clades. The second, GC content difference filtering (absDiffGC), targets loci with reduced base composition bias. These filters are designed to enhance both signal-to-noise ratio and computational efficiency. Our objectives are twofold: (1) to test the monophyly of Squalomorphii and clarify its internal relationships, and (2) to reassess the phylogenetic position of Batoidea relative to Selachii. To further assess the broader utility of the novel metric-based filtering method (absRatioLen), we applied it to a well-studied but contentious vertebrate phylogeny, the sister-group relationship to tetrapods. This case study demonstrates the method’s applicability beyond Chondrichthyes. Overall, our findings highlight the potential of assumption-free filtering approaches to clarify unresolved relationships in the tree of life.

## Methods

### Target Enrichment and Genome Data Retrievement

We applied a targeted gene enrichment method (Li et al., 2012) on 21 chondrichthyan species spanning different orders. This approach was prioritized over whole-genome sequencing due to its higher efficiency, particularly given the exceptionally large genome sizes characteristic of chondrichthyans. Six genomes were compiled (last accessed: 5 January 2019; see Supplementary Table S1) and two well-assembled and annotated genomes, elephant shark (*Callorhinchus milii*) and whale shark (*Rhincodon typus*) were selected as query sequences for baits design. The EvolMarkers (Li et al., 2010; Li et al., 2012) pipeline was used to design baits for capturing chondrichthyan genes. To facilitate BLAST searches, the Perl script, indexandformat.pl was used to construct a genomic index database. Single-copy coding sequences (CDS) longer than 120 bp were extracted from the query species based on gff annotations using singlelarge.pl. Between-genome searches were performed with blastgenomes.pl. Only single-copy one-to-one CDS were retained through using findmarkers.pl and parsed out as potential bait sequences. In total, 24,827 CDS markers were kept for baits design. Further details on the CDS marker design process are illustrated in Supplementary Figure S1. Biotinylated RNA baits were synthesized by Daicel Arbor Biosciences (Ann Arbor, MI, USA). Each bait was designed to be 120 bp in length with a 60 bp tiling overlap between baits allowing 2x complete coverage of each CDS (2x tiling) to increase on-target rate from specific genomic regions (Jiménez-Mena et al., 2022; C. Li et al., 2013). Detailed information on the selected markers and the final bait sequences are provided in the Supplementary Appendix1.

The target enrichment process is described briefly as follows. Genomic DNA was extracted using the Ezup Column Animal Genomic DNA Purification Kit (Sangon, Shanghai, China) following the manufacturer’s instructions and sheared to approximately 250 bp using a Covaris Focused-ultrasonicator M220 (Woburn, Massachusetts, USA). Multiplexed target enrichment sequencing libraries were prepared following the protocol of Meyer and Kircher (2010) with slight modifications (Wang et al., 2022). Gene capture was conducted following the manual of the myBaits Target Sequencing System (Daicel Arbor Biosciences), incorporating modifications as described by Li et al. (2013). The enriched target sequences were pooled at equimolar concentrations and sequenced on an Illumina NovaSeq 6000 (Illumina, Inc, San Diego, CA).

To augment our dataset, the sequence read archive (SRA) data for 97 species (100 samples) were retrieved from NCBI (last accessed: December 23, 2023), as well as 37 available chondrichthyan genomes were downloaded from NCBI, with only the most well-curated genome per species retained (last accessed: June 1, 2024). No genomes from Pristiophoriformes or Echinorhiniformes were found. In total, sequence data from 158 samples across 139 species were compiled to ensure representation of at least one species from each chondrichthyan order (Table 1).

**Table 1.**
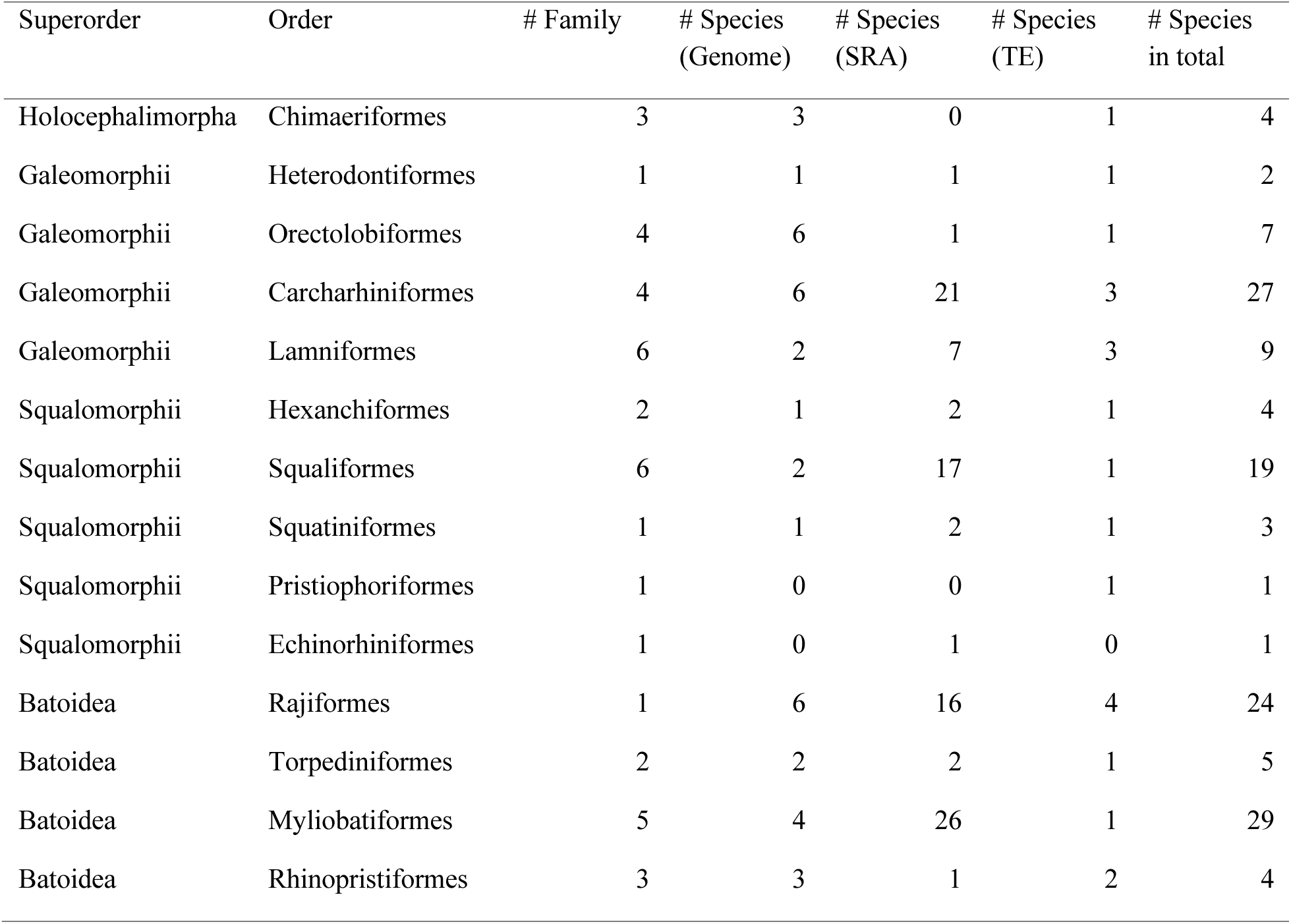
Taxon sampling showing the number of families covered in each chondrichthyan order and the number of species covered by different sequence data resources.

### Target CDS Retrieval and Assembly

Under the Assexon pipeline (Yuan et al., 2019), target genes from the gene capture and SRA data were assembled using the 24,827 developed CDS markers as reference. Target enrichment data was trimmed first to exclude low quality bases and adaptors using trim_galore v0.6.6 (https://www.bioinformatics.babraham.ac.uk/projects/trim_galore/) and Cutadapt v4.1 (Martin, 2011). Orthologues CDS and their flanking sequences were assembled using assemble.pl. The script assemble.pl in Assexon (Yuan et al., 2019) is a wrapper including 1) removal of PCR duplications; 2) parsing of reads to homologous target CDS; 3) assembly of parsed reads; 4) further assembly and retrieval of the best contig per locus; 5) exclusion of potential paralogs following the reciprocal blast strategy. For genome data, orthologues to reference CDS were retrieved using get_orthologues.pl. Information about the initial assemblies (158 samples/139 species/24,827 CDS) can be found in Supplementary Table S2.

### Data Cleanup and Preparation of Data Matrices

The workflow for sequence matrix preparation, phylogenomic analyses, and data filtration is summarized in Figure 2. To resolve ordinal-level relationships within Chondrichthyes, we retained only coding DNA sequences (CDS) present in at least one representative per order using a custom Perl script (decidata.pl; Supplementary Appendix1). To minimize stochastic errors from missing data, we applied the pick_taxa.pl script from the Assexon pipeline, retaining only samples with >3,000 gene loci (final dataset: 98 samples, 85 taxa, 9,663 CDS; Fig. 2). DNA sequences were translated to proteins, aligned with MAFFT v7.520 (Katoh & Standley, 2013), and reverse-translated to DNA using the Assexon pipeline script mafft_aln.pl (Yuan et al., 2019). Codon-based alignments were trimmed with ClipKIT v2.3.0 (Steenwyk et al., 2020) in default “smart-gap” mode to remove uninformative sites and gaps. To address partial taxon coverage bias (Steenwyk et al., 2023), a 70% sample occupancy threshold was enforced, requiring each gene to have ≥69 sequences. This yielded a final nucleotide matrix of 4,738 genes (Fig. 2).

**Fig. 2.**
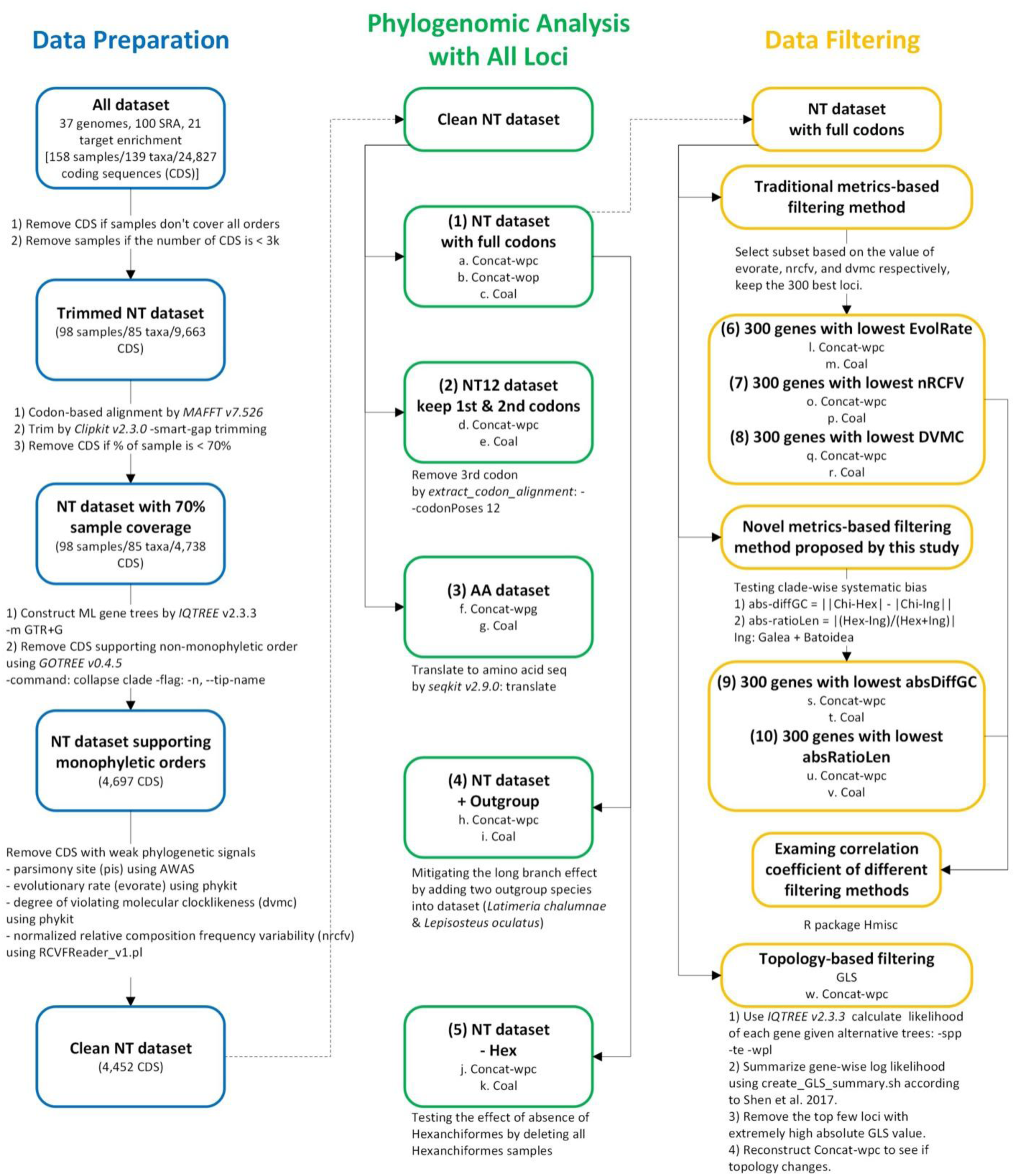
Schematic workflow of the data analyses, divided into three modules: (1) data preparation (sequence alignment, quality control), (2) phylogenomic analyses with all loci (concatenation-based [Concat] vs. coalescent-based [Coal] methods) on different datasets, and (3) data filtering (metric-based vs. topology-based). Partitioning strategies include without partition (*wop*), with partition by codon (*wpc*), and with partition by gene (*wpg*) only applied to amino acid data. Solid arrows indicate sequential steps within each module. Dotted arrows refer to connections between modules.

To identify and exclude erroneous data potentially caused by mislabeling, contamination, misassembly etc., we implemented a two-step filtering approach. First, maximum likelihood (ML) trees were reconstructed for all prefiltered 4,738 loci using IQ-TREE v2.3.3 (Minh, Schmidt, et al., 2020) under the GTR+G substitution model. Next, Gotree v0.4.5 (Lemoine & Gascuel, 2021) was employed to flag CDS non-monophyletic at ordinal-level using the collapse clade command, which replaces designated clades with tip names if they meet monophyly criteria. This generates warnings/errors for non-monophyletic groupings, enabling systematic identification of discordant loci. This method offers distinct advantages over alternatives: constraint-based approaches (e.g., Assexon pipeline; Yuan et al., 2019) rely on statistical comparisons between constrained and unconstrained topologies, which may overlook lineage-specific discordance. By directly testing monophyly without a priori topological assumptions on other relationships, our pipeline ensures robust filtering of gene trees incongruent with ordinal-level classification in Chondrichthyes.

To identify and remove CDS with weak phylogenetic signals, we analyzed sequence alignments using parsimony-informative sites (PIS) calculated with AMAS (Borowiec, 2016), and excluded coding sequences with fewer than 13 PIS. Additionally, we evaluated normalized relative composition frequency variability (nRCFV) (Fleming & Struck, 2023), evolutionary rate (EvolRate) (Telford et al., 2014) and the degree of violation of a molecular clock (DVMC) (Liu et al., 2017) for each gene tree using PhyKIT v1.19.8 (Steenwyk et al., 2021). Outlier loci defined as values exceeding 1.5 times the interquartile range for nRCFV, EvolRate, and DVMC were excluded from subsequent analyses. These criteria were not designed to prioritize phylogenetically informative loci but to eliminate potential artifacts arising from homoplasy or assembly errors. The resulting filtered dataset, named “Clean NT”, was retained for downstream analyses (Fig. 2; Dataset 1).

To mitigate effect of substitution saturation at third codon positions (Breinholt & Kawahara, 2013), we generated a nucleotide matrix (NT12) using the extract-codon-alignment Python tool (Cock et al., 2009) with parameter -codonPoses 12, retaining only first and second codon positions (Fig. 2; Dataset 2). Nucleotide matrices are computationally efficient and accommodate advanced substitution models, while amino acid matrices reduce homoplasy (Kapli et al., 2023). For comparatively deep phylogenetic inference, we translated nucleotide sequences to amino acids (AA matrix; Fig. 2; Dataset 3) using SeqKit v2.9.0 (Shen et al., 2016).

Li et al. (2012) found that base compositional non-stationarity and distance from the root compromised accuracy of estimated relationships within the ingroup of chondrichthyans and biased base composition of hexanchiform species led to erroneous phylogenetic reconstruction. To address the long branch between Chimaeriformes and Elasmobranchii (Li et al., 2012), we expanded outgroup sampling to include the West Indian Ocean coelacanth (*Latimeria chalumnae*) and spotted gar (*Lepisosteus oculatus*), creating the “NT+Outgroup” matrix (Fig. 2; Dataset 4). To test the topological influence of Hexanchiformes, we constructed an “NT−Hex” matrix by excluding hexanchiform samples (Fig. 2; Dataset 5).

### Phylogenomic Analyses with All Loci

Both concatenation-based and coalescent-based approaches were implemented. For the concatenation-based approach, the GTR+G model was applied to the NT dataset (including NT12, “NT+Outgroup”, and “NT-Hex”) with two different partition schemes (without partition and partitioned by codon). The LG+G model was used for the AA dataset, partitioned by gene. The ML tree was reconstructed using IQ-TREE v2.3.3 (Minh et al., 2020) and the topological robustness of each tree was evaluated with 1,000 ultrafast bootstrap replicates (Minh et al., 2013). For the coalescence-based approach, species trees were inferred from individual ML gene trees using ASTRAL v5.7.8 (Mirarab et al., 2014), which accounts for incomplete lineage sorting (ILS). Topological robustness was evaluated via local posterior probabilities (LPP). Gene concordance factors (gCF) and site concordance factors (sCF) were calculated for the NT dataset using IQ-TREE (Minh et al., 2020). Phylogenetic incongruence among trees inferred from various datasets and methodological approaches was visualized as a heatmap across intra- and inter-superorder levels.

### Data Filtration Based on Extant Methods

Metric-based and topology-based filtering methods represent two principal strategies for investigating phylogenetic incongruence (Steenwyk et al., 2023). In metric-based filtering, systematic errors, such as compositional heterogeneity and substitution rate variation, were evaluated using three conventional phylogenetic signal metrics: nRCFV, EvolRate, and DVMC. Genes exhibiting robust phylogenetic signals (characterized by low values across all three metrics) were prioritized for downstream phylogenetic reconstruction. The first metric, nRCFV, quantifies compositional heterogeneity and was calculated using the RCFVReader_v1.pl script. This method accounts for alignment length, taxon number, and character state composition (Fleming & Struck, 2023). The remaining metrics, EvolRate and DVMC, serve as proxies for substitution rate variation and clock-likeness. These were computed with PhyKIT v1.19.8 (Steenwyk et al., 2021) based on inferred gene trees. EvolRate reflects the total tree length normalized by the number of terminal taxa (Telford et al., 2014), while DVMC measures the standard deviation of root-to-tip distances across species (Liu et al., 2017). For each metric, the 300 genes with the lowest values were retained for analyses (Fig. 2; Datasets 6–8).

Topology-based filtering methods, such as gene genealogy interrogation (GGI) (Arcila et al., 2017) and gene-wise likelihood scoring (GLS) (Shen et al., 2017), require predefined phylogenetic hypotheses as constraints to calculate per-gene log-likelihood scores. This enables the identification of loci supporting specific topologies. Here, we implemented GLS to compare it with metric-based filtering methods. Two hypothetical topologies, derived from the most frequent polytomies (while preserving ordinal-level monophyly) in ML and multispecies coalescent (MSC) results—were used as constraints. Per-gene likelihood scores were computed with IQ-TREE v2.3.3 (Minh et al., 2020) using the -spp, -te, and -wpl parameters. Following Shen et al. (2017), we calculated the difference in likelihood scores (ΔGLS) for each gene. Genes with extreme ΔGLS value were discarded as outliers based on scatterplot. ML and MSC analyses were rerun using the outlier-free dataset. The impact of outlier loci on phylogenetic resolution was assessed by examining whether the sign of overall ΔGLS is flipped, and visualizing the topologies reconstructed before and after filtering. This approach aligns with established practices for resolving contentious phylogenies (Ballesteros & Sharma, 2019; Hughes et al., 2023; Li et al., 2021; Shen et al., 2017).

### Novel Data Filtering Methods Proposed in This Study

While traditional phylogenetic signal metrics (e.g., nRCFV, EvolRate, DVMC) evaluate biases at the locus-wide level, they overlook heterogeneity introduced by individual taxa or subgroups within a locus. Building on Li et al. (2012)’s findings that differences in root branch lengths and base composition between outgroup and ingroup taxa can compromise phylogenetic accuracy, we introduce two novel fine-scale phylogenetic signal metrics to detect deviations among subgroups. The first metric, absolute difference in GC content between focal clades (absDiffGC) (Equation 1), serves as a proxy for compositional heterogeneity, which is a known source of systematic error in phylogenetic inference. The second metric, absolute ratio of branch length (absRatioLen) between focal clades (Equation 2), which captures rate asymmetry among groups and addresses ingroup-outgroup imbalance.

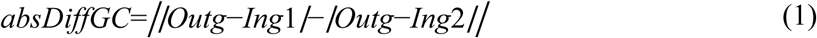

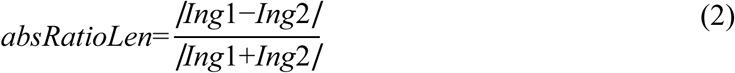

The focal clades can be identified by comparing branch lengths and topologies reconstructed from the unfiltered dataset. For both metrics, genes were partitioned into three groups, one outgroup and two ingroups (with one designated as the focal group). Preliminary phylogenetic analyses with all loci revealed conflicts in the placement and monophyly of Squalomorphii (Figs 3 and 4). Maximum likelihood analyses indicated that Selachii are paraphyletic, with Galeomorphii clustering with Batoidea and Squalomorphii positioned basally (Fig. 3). Within this arrangement, Hexanchiformes emerged as the sister group to the remaining Squalomorphii. Analysis of the “NT-Hex” matrix—which excluded Hexanchiformes—underscored the significant impact of Hexanchiformes heterogeneity on elasmobranch topology (Fig. 4 and Supplementary Fig. S2-S3). The “NT-Outgroup” matrix, incorporating two actinopterygian species, highlighted the recalcitrant influence of outgroup (Chimaeriformes) distance (Supplementary Fig. S4-S5).

**Fig. 3.**
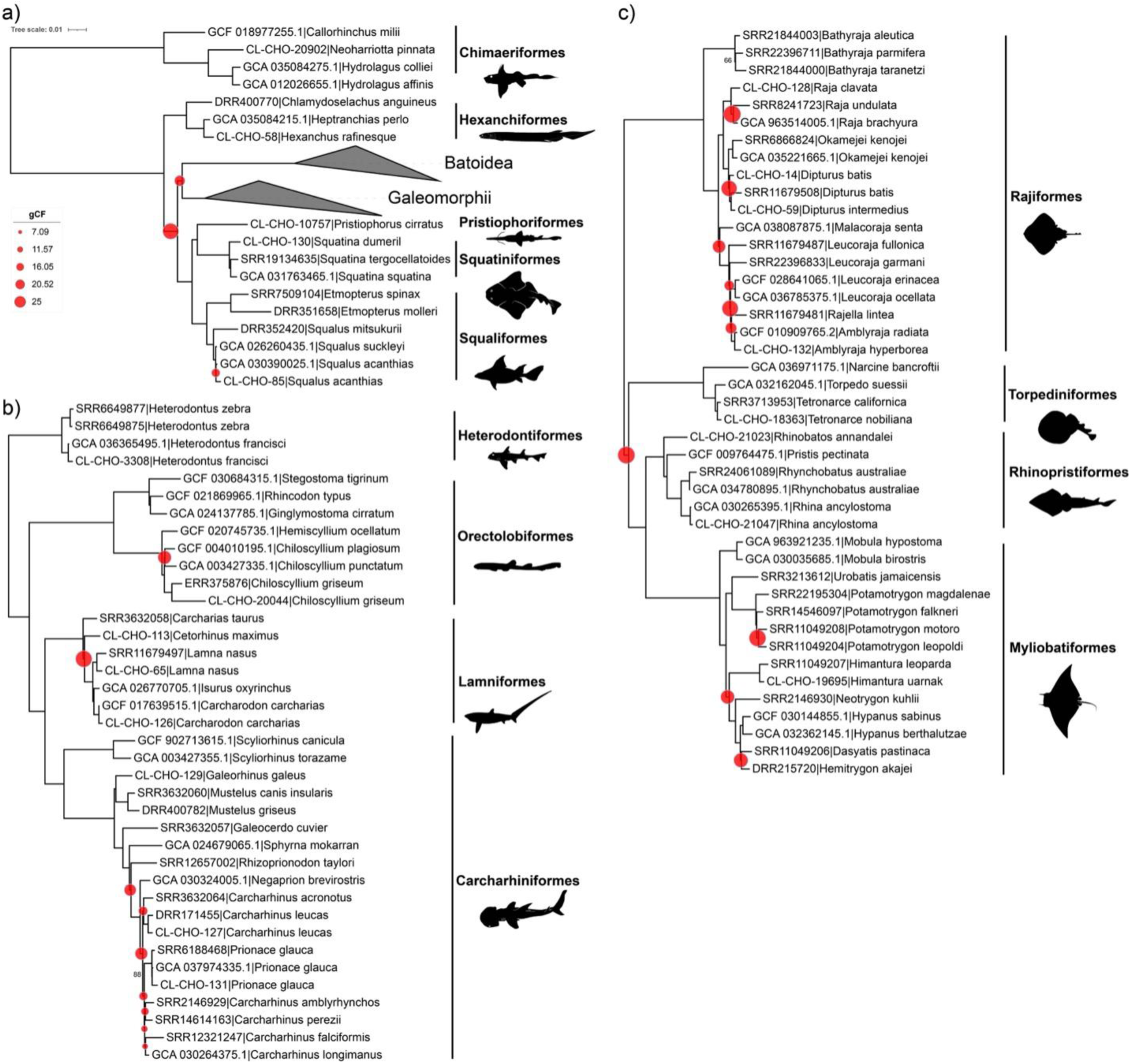
Maximum likelihood phylogeny reconstructed from NT dataset 1 using IQ-TREE under a codon-partitioned scheme. The tree was visualized in iTOL (Letunic and Bork, 2004), with bootstrap support values below 90% labeled. Ordinal level nodes exhibiting gene concordance factors (gCF) below 25% are marked by red proportional circles, scaled to reflect gCF magnitude. Tip names are labeled with species name followed by accession numbers. Silhouettes are taken from the public domain Phylopic.org. (a) Overview of chondrichthyan phylogeny with collapsed Galeomorphii and Batoidea. (b) Expanded view of Galeomorphii. (c) Expanded view of Batoidea.

**Fig. 4.**
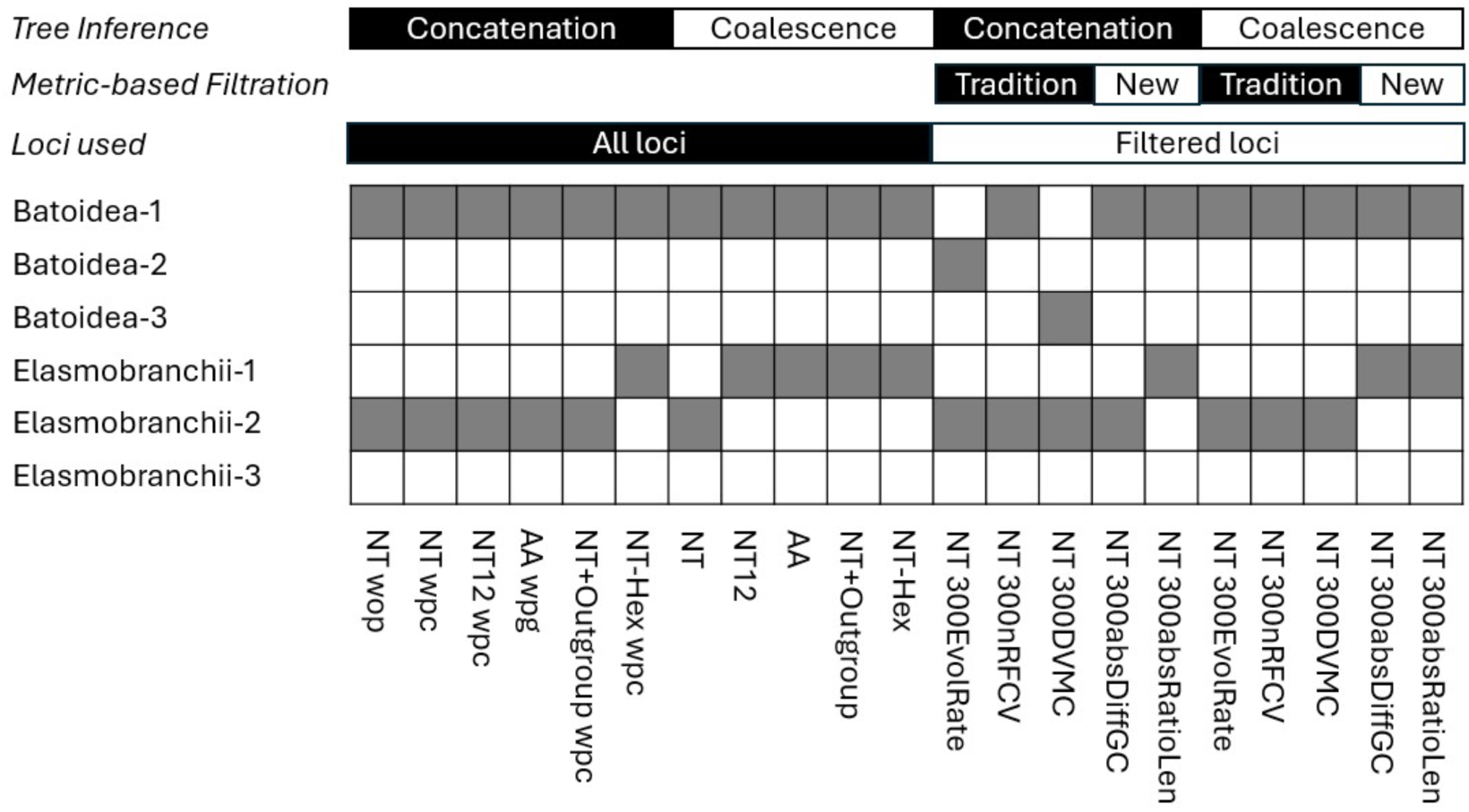
Summary of topological support for controversial Elasmobranchii and Batoidea relationships across 21 phylogenetic reconstructions, incorporating various analytical regimes: inference frameworks (concatenation [partitioning schemes: unpartitioned *wop*, codon-partitioned *wpc*, gene-partitioned *wpg*] vs. coalescent-based methods), data types (full nucleotide sequences, 1^st^+2^nd^ codon positions, amino acid translations), taxon sampling strategies (expanded outgroups [+2 taxa] vs. Hexanchiformes exclusion), and phylogenetic signal filtering approaches (traditional metrics: evolutionary rate—EvolRate, relative composition frequency variation— nRCFV, site concordance—DVMC; novel metrics: GC-content divergence—absDiffGC and branch-length ratio—absRatioLen).

To implement our proposed metrics, we defined the taxonomic groupings as follows: Chimaeriformes served as the outgroup (“Outg”), Hexanchiformes as the first ingroup (“Ing1”), and Galeomorphii together with Batoidea as the second ingroup (“Ing2”) (Fig. 5 and Supplementary Fig. S2-S5). In theory, an absDiffGC value near zero indicates minimal divergence in base composition between Hexanchiformes and the other ingroups relative to Chimaeriformes, reflecting compositional homogeneity. Similarly, an absRatioLen value approaching zero suggests that branch lengths of Hexanchiformes are comparable to the average ingroup branch length. However, because branch length variation is influenced by both base composition and substitution rate heterogeneity, even loci with similar GC content may display divergent branch lengths.

**Fig. 5.**
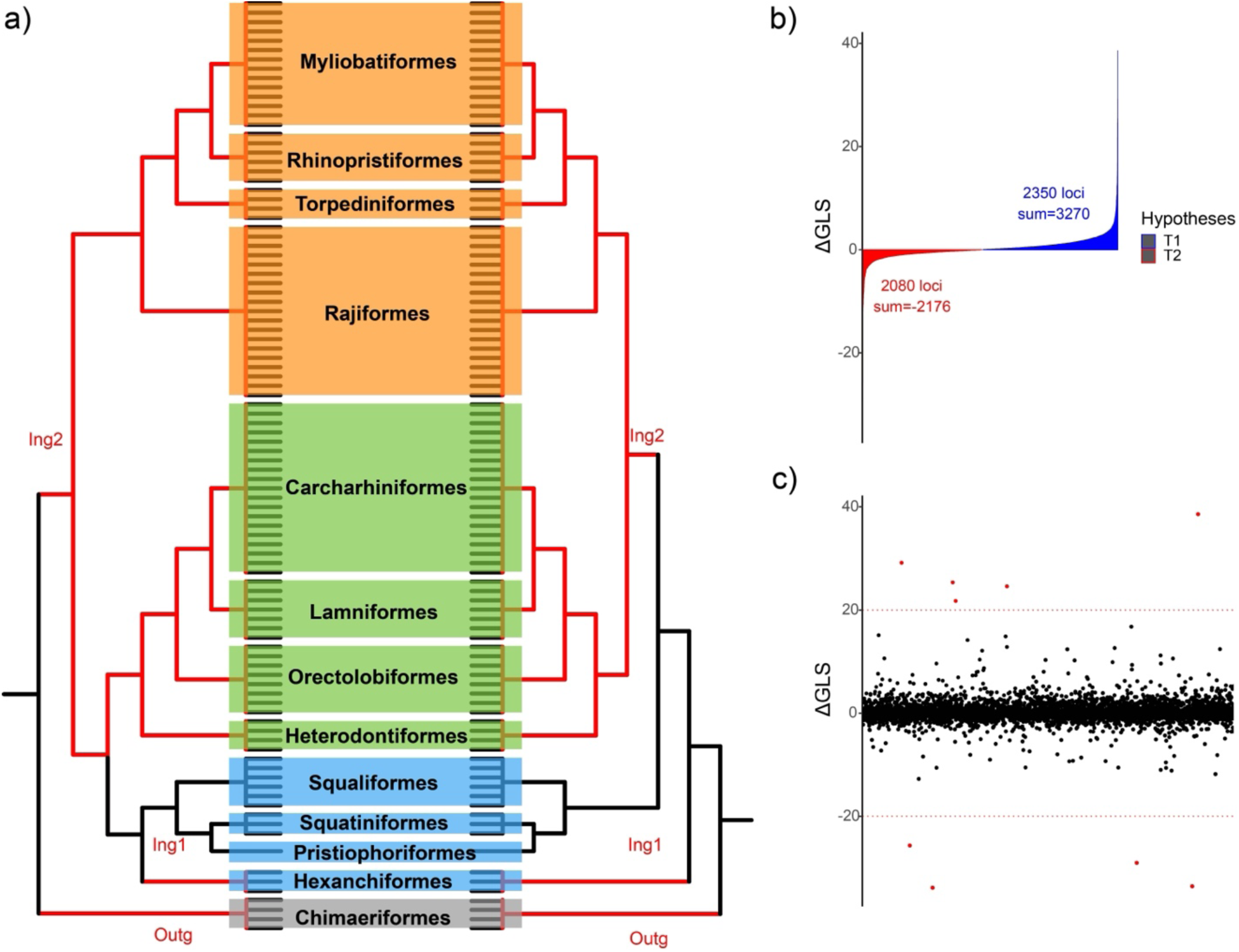
(a) Alternative hypotheses on chondrichthyan phylogenetic relationship: left (Elasmobranchii-1) resolves Selachii as monophyletic, and right(Elasmobranchii-2) exhibits a paraphyletic Selachii—a pattern frequently observed in concatenation-based trees. Color shading: orange denotes the Batoidea clade, green denotes the Galeomorphii clade, and blue denotes the Squalomorphii clade. The red branches highlight the clades selected for novel metrics measurements, with labels on branches specifying subgroups for calculating the metrics. (b) Bar plot comparing the number of loci supporting competing topologies: T1 (blue; 2,350 loci, cumulative ΔGLS = 3,270.4, max = 38.59) versus T2 (red; 2,080 loci, cumulative ΔGLS = - 2,176, min = -33.86). The global mean ΔGLS across all loci is 0.25. (c) Scatter plot of locus-specific ΔGLS values, with outliers defined as values beyond ±20 (indicated by red dashed lines). Nine outlier loci (red dots) were identified.

To identify informative loci which are less affected by systematic errors, we retained the 300 genes (<10%) with the lowest absDiffGC (Fig. 2; Dataset 9) and absRatioLen (Fig. 2; Dataset 10), respectively, constructing two distinct data matrices. These datasets were analyzed using standard phylogenomic pipelines to infer evolutionary relationships. To evaluate the correlation between phylogenetic signals (the values of the five metric) from novel (localized, lineage-specific) and traditional (global average) metrics, we calculated Pearson correlation coefficients for each locus in the NT dataset 1 and assessed their statistical significance using the *corrplot & Hmisc* package in R (Jr, 2025; Wei & Simko, 2024). To examine the composition of sub-matrices selected based on the most informative phylogenetic signals, we computed and visualized their intersections using the R package *ggupset* (Ahlmann-Eltze, 2025). The size and direction (positive or negative) of ΔGLS were used to quantify topological support. To assess trends between ΔGLS and each metrics, metrics values were ordered ascendingly, and local polynomial regression was fitted to plot the trend line using R in loess function.

### Resolving a Classic Vertebrate Phylogenetic Conflict Using Novel Data Filtering

To assess the broader utility of our metric-based filtering approach, we applied it to the long-debated relationships among lungfish, coelacanths, and tetrapods. While mounting evidence supports lungfish as the sister group to tetrapods (Irisarri et al., 2017; Irisarri & Meyer, 2016), this relationship remains contested. We analyzed 4,593 amino acid alignments from Irisarri et al. (2017), reducing the dataset to 24 representative taxa for computational efficiency. Maximum likelihood analyses were conducted in IQ-TREE using a gene-partitioned scheme with the LG+G model.

Following Takezaki and Nishihara’s (2017) protocol, we evaluated five outgroup configurations: (1) Chondrichthyes + ray-finned fishes (Holostei + Teleostei), (2) Chondrichthyes only, (3) ray-finned fishes only, (4) Chondrichthyes + Teleostei, and (5) Teleostei only. Given that closely related outgroups may bias phylogenetic inference (Takezaki & Nishihara, 2017), we focused our absRatioLen filtering analysis on configuration 4 (Chondrichthyes + Teleostei) to minimize potential artifacts. The filtered loci were then used to reconstruct the final phylogeny and evaluate topological stability.

## Results

### Data Compilation, Quality Control, and Cleanup

A total of 24,827 single-copy coding sequences (CDS) were identified from six chondrichthyan genomes for RNA bait development. The complete chondrichthyan bait sequences are publicly accessible in the repository Rhincodon_typus.dna.fasta (Supplementary Appendix1). These baits were subsequently applied in target enrichment experiments across 21 chondrichthyan samples. Altogether, we compiled sequence data from 158 samples representing 139 species, including 37 whole genomes, 100 SRA datasets, and 21 target enrichment (TE). Trimmed sequence data of target enrichment samples are available under NCBI Bioproject PRJNA1232202. Assembled and retrieved target sequences are detailed in Supplementary Table S2. The single echinorhiniform sample yielded only 430 CDS post-assembly, far below the 3,000-gene threshold for initial filtering. However, as it represents the sole member of this order and our study aims to resolve ordinal-level phylogeny in Chondrichthyes, so we created a separate dataset including this sample to infer its phylogenetic position (see Discussion).

During data filtering, we removed 4,925 CDS that exceeded 30% missing data across samples, 41 CDS that yielded non-monophyly at the ordinal level, and 8 CDS with too few parsimony-informative sites (PIS <13), retaining 4,689 CDS. This dataset was further refined by calculating the interquartile range (IQR) for three phylogenetic signal metrics and removing outliers (values >1.5×IQR), resulting in a final nucleotide sequence matrix of 4,452 CDS (Supplementary Fig. S6; designated as NT dataset 1). This matrix comprises 52,855,656 sites, with assembly details in Supplementary Table S3. The full nucleotide sequence matrix was transformed into nine additional matrices (dataset 2-10 in Fig. 2) by applying various criteria: extracting only first and second codon positions, translating into amino acids, adding outgroups, removing Hexanchiformes, and selecting the 300 most informative loci based on three traditional and two novel metrics. These nine matrices (Supplementary Table S4) span 300–4,452 loci and 95–100 samples, containing 3,449,298–54,580,527 characters. Missing data (alignment gap) percentages ranged from 1.016% to 3.183%. The distribution of six phylogenetic signal metrics for NT dataset 1 is illustrated in Supplementary Figure S7.

### Phylogenetic Reconstruction and Topological Discordance Based on All Loci

Five matrices (dataset 1-5: NT, all nucleotides; NT12, nucleotide data with 1st&2nd codons only; AA, amino acid sequences; NT+Outgroup, nucleotide data with two extra outgroups; NT-Hex, nucleotide data without hexanchiforms; see Fig. 2) containing all 4,452 filtered loci were analyzed to reconstruct chondrichthyan phylogenies using two tree-inference methods (concatenation vs. coalescent) and three partitioning strategies (without partition, partitioned by codon, and partitioned by gene; Fig. 2). The concatenation-based analysis of NT dataset 1 with codon-partitioning revealed a topology (Fig. 3) characterized by two notable features: the non-monophyly of Squalomorphii and the sister-group relationship between Galeomorphii and Batoidea, consistent with the findings of Li et al. (2012) (Fig. 1b). Bootstrap support values reached almost 100% for all ordinal divergences (Supplementary Fig. S4). However, three critical ordinal-level nodes exhibited gene concordance factor (gCF) values below 25% (Fig. 3). These included the divergence of Hexanchiformes (e.g., cow shark) from other elasmobranchs (sharks, skates and rays), the split between Galeomorphii (modern sharks) and Batoidea (rays and skates), and the differentiation of Torpediniformes (e.g., electric rays) from Myliobatiformes (e.g., stingrays) and Rhinopristiformes (e.g., guitarfish). We observed intra-ordinal discordance within several groups, including Myliobatiformes and Rhinopristiformes (Batoidea), as well as Lamniformes, Orectolobiformes, and Squaliformes (Selachimorpha). Notably, Rajiformes and Carcharhiniformes exhibited heightened conflict, each containing more than five nodes with low gCF values.

We generated in total eleven different phylogenies (6 from concatenation method, 5 from coalescence method) from the full dataset (4,452-loci), revealing systematic discordance across analytical approaches and the effect of adding extra outgroups or excluding Hexanchiformes (Fig. 4 Left). All trees supported the Batoidea-1 topology, which positions the Rajiformes basally and recovers the Torpediniformes as the sister group to the Rhinopristiformes and Myliobatiformes (Fig. 1d). For contentious relationships within Elasmobranchii, concatenation-based methods predominantly supported the Elasmobranchii-2 configuration, with the exception of analyses using the NT-Hex matrix. Coalescent-based methods favored the Elasmobranchii-1 topology for NT12, AA, NT+Outgroup, and NT-Hex, whereas the NT dataset supports Elasmobranchii-2. These results highlight methodological influences on phylogenetic resolution and underscore persistent uncertainties in the determination of chondrichthyan evolutionary relationships.

### Phylogenetic Reconstructions Using Filtered Datasets

#### Traditional Metric-Based Filtering Analyses

Three matrices derived from NT dataset 1 through traditional metric-based filtering (nRCFV, DVMC, and EvolRate) were used to reconstruct six phylogenies with distinct inference methods. All resulting trees consistently supported the Elasmobranchii-2 topology (Fig. 4). Concatenation-based methods exhibited variability in Batoidea topology resolution: only the phylogeny from the “300nRCFV” matrix recovered the Batoidea-1 topology, while the other two matrices produced alternative Batoidea hypotheses. In contrast, coalescent-based methods uniformly resolved the Batoidea-1 topology across all analyses (Fig. 4 Right).

#### Topology-Based Filtering Analyses

To assess these competing hypotheses (Fig. 5a), gene-wise likelihood score (GLS) calculations were performed across all 4,452 loci under constrained topologies. Of these, 2,350 loci favored T1 and 2,080 supported T2. The total ΔGLS is +1,094, which overall suggests a preference for the T1 topology. Extreme ΔGLS values were comparable between hypotheses, with a maximum ΔGLS of +38.59 for T1 and a minimum ΔGLS of –33.86 for T2 (Fig. 5b). Nine loci were identified as extreme outliers through a scatter plot (Fig. 5c). Exclusion of these outliers reduced the total ΔGLS from +1,094 to +1,077 but still favored the T1 topology. The favored topology only shifts from T1 to T2 when the ΔGLS value becomes negative. Phylogenetic reconstructions using the pruned matrix (4,443 loci) under both concatenated and coalescent frameworks produced topologies identical to those derived from the full dataset (Supplementary Fig. S8-S11).

### Results of Novel Localized Metric-Based Filtering

In this study, Chimaeriformes was treated as the outgroup (Fig. 3). Hexanchiformes was recovered either within a monophyletic Squalomorphii or as part of paraphyletic Squalomorphii (Fig. 4a). It is notable that in the paraphyletic Squalomorphii, all members except Hexanchiformes formed a monophyletic group, suggesting that Hexanchiformes may be pulling them out from their sister relationship with Galeomorphii. The branch length of the Hexanchiformes to the root is short (ing1), whereas the branch lengths to the families of Galeomorphii and Batoidea are much longer (ing2). Thus, we define the Hexanchiformes as ingroup 1 and the Galeomorphii and Batoidea families as ingroup 2 and calculate metrics using our newly proposed methods accordingly.

Phylogenetic analyses using datasets filtered by our novel localized metrics consistently recovered the Batoidea-1 topology across both concatenation- and coalescent-based methods (Fig. 4). However, resolution of Elasmobranchii relationships varied between matrices. The 300absRatioLen matrix uniformly supported Elasmobranchii-1 under all inference frameworks (Fig. 4; Fig. 6), while the 300absDiffGC matrix resolved Elasmobranchii-1 in coalescent-based analyses but shifted to Elasmobranchii-2 in concatenation-based reconstructions (Fig. 4). Notably, despite its support for Elasmobranchii-1, the 300absRatioLen matrix revealed an atypical arrangement within Squalomorphii, positioning Squatiniformes as sister to Squaliformes + Pristiophoriformes (Supplementary Fig. S12)—a departure from the conventional topology placing Squaliformes as sister to Squatiniformes + Pristiophoriformes (Supplementary Fig. S13).

**Fig. 6.**
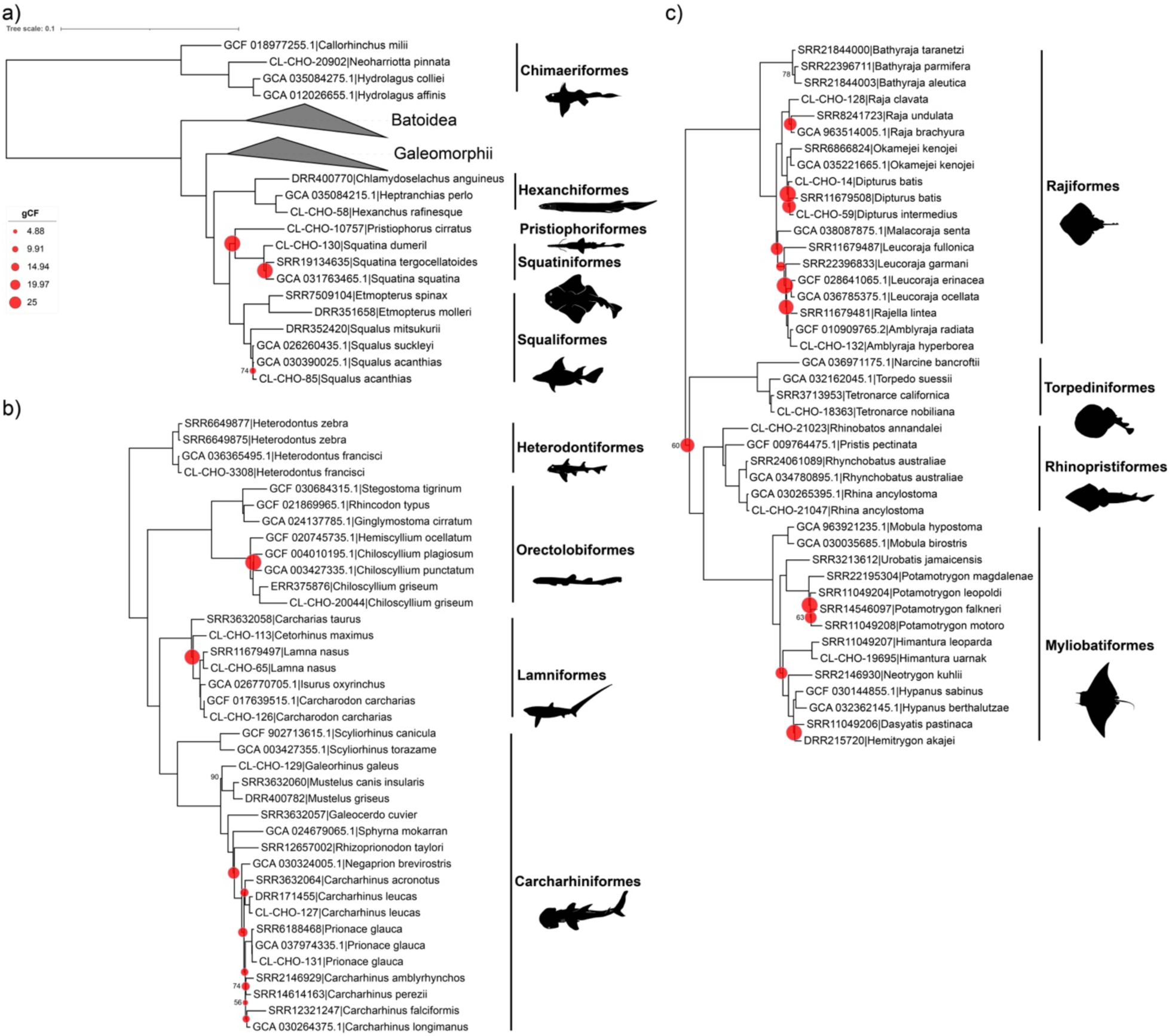
Maximum likelihood phylogeny reconstructed from NT300absRatioLen dataset using IQ-TREE under a codon-partitioned scheme. The tree was visualized in iTOL (Letunic and Bork, 2004), with bootstrap support values below 90% labeled. Tip names are labeled with species name followed by accession numbers. Four superorders are shaded by color. Silhouettes are taken from public domain of Phylopic.org. (a) Overview of Chondrichthyan phylogeny with collapsed Galeomorphii and Batoidea. (b) Expanded view of Galeomorphii. (c) Expanded view of Batoidea.

### Correlations between Phylogenetic Signal Metric

Pearson correlation analysis of phylogenetic signal metrics in the cleaned NT dataset 1 (α = 0.05) revealed distinct patterns of interdependence (Supplementary Tables S6–S7). The absRatioLen metric exhibited no significant correlations with other metrics except for a small but negative association with EvolRate (p<0.05, Fig. 7a). In contrast, the remaining four metrics—DVMC, nRCFV, EvolRate, and absDiffGC—showed significant positive correlations. We observed strong inter-metric relationships (Pearson’s *r* > 0.37) among DVMC, nRCFV, and EvolRate, while absDiffGC displayed moderate but significant correlations with these traditional metrics. Additionally, only absRatioLen displayed a consistent directional relationship with ΔGLS across datasets, a pattern not observed for the other metrics (Supplementary Fig. S31). (Supplementary Figure S31).

**Fig. 7.**
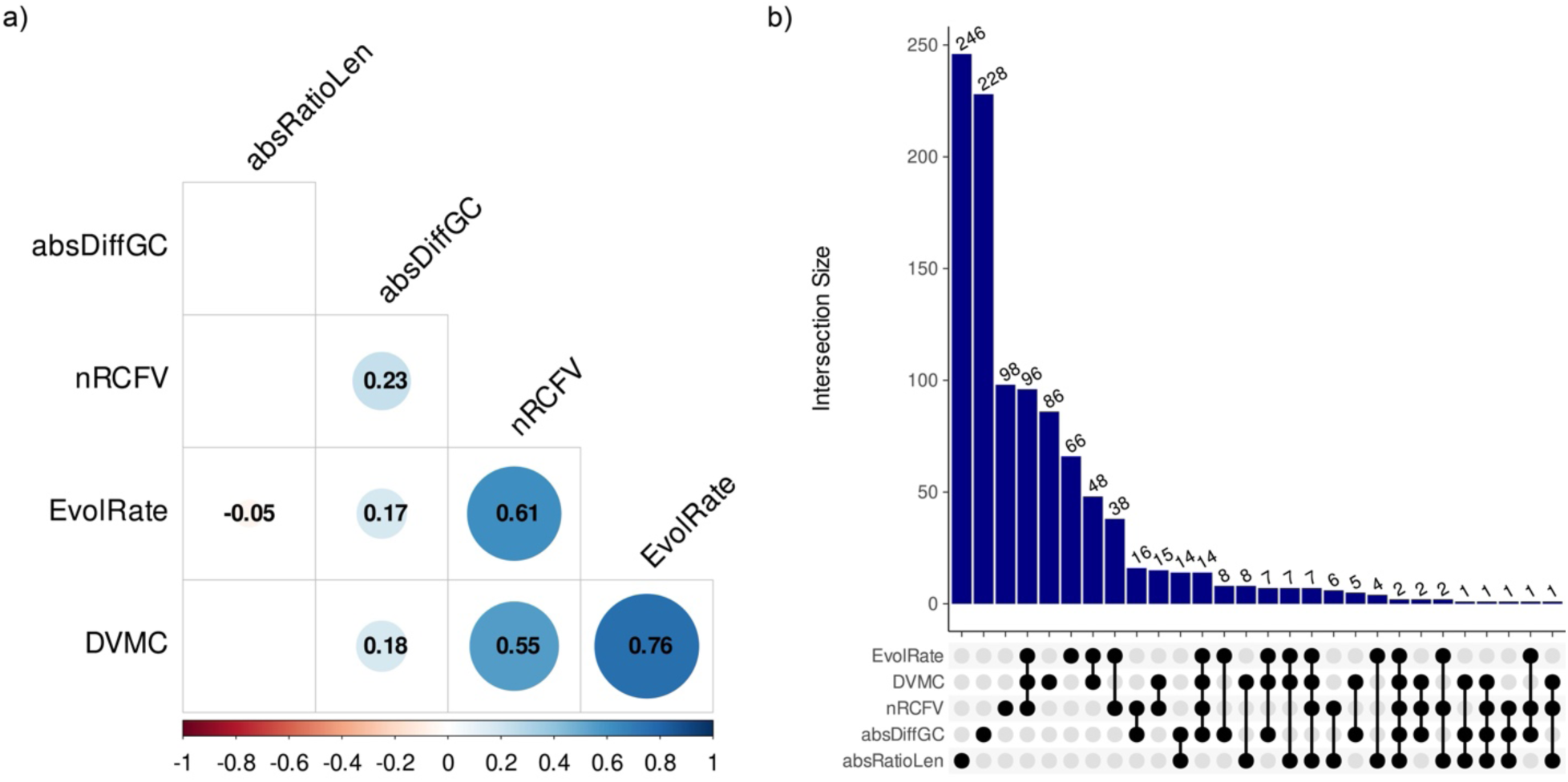
Pairwise correlations among phylogenetic signal metrics and intersections among the top 300 most informative CDS. (a) Correlation matrix: Metrics analyzed include normalized relative composition frequency variation (nRCFV), evolutionary rate (EvolRate), degree of molecular clock violation (DVMC), absolute GC-content divergence (absDiffGC), and absolute branch-length ratio (absRatioLen). Each cell with statistically significant correlations is annotated with the corresponding correlation coefficient. Blue and red circles represent significant positive and negative correlations, respectively (p < 0.05), with both circle size and color intensity proportional to the magnitude of Pearson’s *r* (numbers in the circles). b) Intersection analysis: The vertical bar plot displays the number of CDS shared among different metrics, which were indicated by the black-filled dots under the y-axis.

When we examined the overlap of the top 300 loci selected by each metric (Fig. 7b), we found that the absRatioLen and absDiffGC metrics shared few loci with other metrics but contained a large number of unique loci, accounting for 83% (246 loci) and 76% (228 loci), respectively. However, traditional metrics (nRCFV, DVMC, EvolRate) had fewer exclusive loci (<100 each), with 96 loci common to all three. Notably, just two loci were shared universally across all metrics. These results underscore the distinctiveness of the signals selected by using the localized versus traditional filtering approaches.

### Untangling a Classic Vertebrate Phylogenetic Dispute

With Chondrichthyes + ray-finned fishes, Chondrichthyes only, or Holostei + Teleostei as outgroups, lungfish were consistently recovered as the closest relatives of tetrapods (Supplementary Figure S33 A-C), which was demonstrated in Takezaki and Nishihara (2017) about the outgroup selection effect. In contrast, analyses with Teleostei only or Chondrichthyes + Teleostei as outgroups supported a sister relationship between lungfish and coelacanths (Supplementary Figure S33 D-E). To investigate whether tetrapod branch lengths influenced lungfish and coelacanth placement, lungfish were designated as ingroup1 and tetrapods as ingroup2. We then applied the absRatioLen metric-based filtering method, selecting the top 300 most informative loci for ML reconstruction. The filtered dataset recovered lungfish as the closest relatives of tetrapods (Supplementary Figure S33 F), even when Teleostei were used as the outgroup, contradicting to the findings of Takezaki and Nishihara (2017).

## Discussion

### Traditional Approaches to Mitigate Systematic Error

Phylogenomic analyses commonly employ one of two methodological frameworks: either concatenated supermatrix approaches or coalescent-based gene-tree methods. The supermatrix method amplifies strong phylogenetic signals (Smith et al., 2020) but risks magnifying systematic biases, such as substitution model misspecification, ortholog misassignment, and rooting errors (Li et al., 2012; Steenwyk et al., 2023), which can exacerbate incongruence as datasets scale. Conversely, coalescent approaches explicitly model biological sources of discordance, such as incomplete lineage sorting (ILS), by integrating gene tree quartets (Edwards et al., 2007; Maddison, 1997). While coalescent methods often yield more biologically plausible species trees, they cannot address problems of systematic errors and their accuracy hinges on the precision of individual gene tree estimations (Springer & Gatesy, 2016).

To mitigate systematic errors in phylogenomic analyses, multiple data filtering criteria have been proposed. These approaches often focus on excluding rapidly evolving sites, such as third codon positions, reducing substitution saturation (Jeffroy et al., 2006) or translating nucleotide sequences into amino acids to lessen the influence of fast-evolving nucleotide bias (Li et al., 2012). Compositional heterogeneity is addressed by prioritizing loci with low Relative Composition Frequency Variability (RCFV) (Zhong et al., 2011), while molecular clock-like genes are selected to align with homogeneous evolutionary models (Kuang et al., 2018; Liu et al., 2017). Collectively, these methods aim to improve phylogenetic accuracy by filtering loci based on compositional stability and evolutionary rate homogeneity.

Empirical results from this study, however, suggest limited efficacy of conventional metrics in enhancing phylogenetic congruence. For example, phylogenies reconstructed from nucleotide datasets excluding third codon positions (NT12) and amino acid datasets (AA) remained incongruent across different inference methods (e.g., concatenation vs. coalescence), despite internal consistency within each methodological framework (Fig. 4). Subsets of markers selected using metrics such as EvolRate, nRCFV, and DVMC exacerbated topological discordance, particularly regarding the contentious Batoidea relationships. This increased discordance likely stems from heightened sampling error due to reduced data volumes. Further compounding these issues is the independent application of filtering criteria without accounting for interdependencies among metrics, coupled with dataset size constraints (100–200 markers) (Arcila et al., 2017; Li et al., 2021).

### Comparison Between Conventional and Novel Metric-Based Filtering Methods

Subsampling genes with strong signal-to-noise ratios is a widely adopted strategy in phylogenomics, particularly for mitigating systematic errors amplified by large datasets (Arcila et al., 2017; Li et al., 2012; Simmons et al., 2016). Traditional metric-based filtering approaches calculate phylogenetic signals, such as base composition or evolutionary rate homogeneity across entire loci. We evaluated three conventional metrics: nRCFV, which quantifies base compositional bias; EvolRate, measuring evolutionary rate heterogeneity; and DVMC, assessing deviations from clock-like evolution (Liu et al., 2017). These metrics were applied globally, averaging signals across all taxa per gene, a common practice that risks overlooking intra-clade heterogeneity. Conversely, our novel metric-based filtering methods focus on problematic clades and evaluate genes on their characteristics at local clades, so they may reveal the true problematic loci causing errors in phylogenetic inference.

In the concatenated tree from unfiltered dataset, Hexanchiformes exhibit notably shorter branch lengths than other elasmobranchs, resulting in their placement at the base of Elasmobranchii and rendering Squalomorphii paraphyletic (Fig. 1). Similar patterns were reported by Li et al. (2012), who demonstrated that deviations from base compositional stationarity and increasing distance from the root can reduce the accuracy of inferred relationships within the ingroup. To address such limitation of clade-specific biases, Smith et al. (2023) refined the traditional PIS metric by scaling each individual’s PIS by the average PIS per locus to help identify samples with low information content. However, this metric is only used to filter out outlier taxa but not “bad” loci, and PIS only reflects variation based on site differences in the alignment. In contrast, branch length provides complex evolutionary signals, capturing both substitution rate and base composition biases.

Correlation analyses of 4,452 loci from the Clean NT dataset (Fig. 2) revealed strong positive correlations among the three traditional metrics. In contrast, absDiffGC showed only weak positive correlations with all metrics except absRatioLen, while absRatioLen exhibited no correlation with other metrics, except only a weak association with EvolRate (Fig. 7a). The superior performance of absRatioLen compared to absDiffGC likely reflects fundamental differences in their underlying biological signals. While absDiffGC measures nucleotide frequency variations alone, absRatioLen incorporates both the number and type of substitutions, thereby capturing deeper evolutionary patterns including homoplasy and substitutional saturation, factors undetectable through GC content analysis. This distinction emphasizes the greater phylogenetic utility of branch length as a more comprehensive evolutionary metric.

When selecting the top 300 loci based on phylogenetic signal, absDiffGC and absRatioLen identified largely distinct gene sets, whereas traditional metrics shared approximately 67% of selected loci (Fig. 7b). This substantial overlap among traditional metrics confirms their interdependence, while the independence of our novel metrics aligns with their distinct correlation patterns. Crucially, phylogenies reconstructed from absRatioLen-selected loci demonstrated superior topological consistency compared to those derived from either absDiffGC or traditional metrics. This enhanced performance likely stems from branch length’s capacity to integrate both base composition effects and substitution rate variation, providing a more complete representation of lineage-specific evolutionary processes.

The effectiveness of absRatioLen is particularly noteworthy given the established value of branch length metrics in detecting systematic errors. These include long-branch attraction (LBA), where rapidly evolving taxa are artifactually grouped (Felsenstein, 1978), and outgroup-induced topological distortions (Lartillot et al., 2007; Li et al., 2012). The robust performance of absRatioLen in resolving deep phylogenetic conflicts reaffirms the importance of branch length-based approaches in phylogenomic analyses.

### Increasing the number of selected loci

To assess the stability of phylogenetic inference under relaxed filtering criteria, we selected the top 500 and 700 most informative loci based on three traditional metrics and two novel metrics. Intersection analyses (Supplementary Fig. S27) showed a similar pattern to the 300-locus datasets: each novel metric retained a high proportion of (60–70%) unique loci, although this proportion decreased slightly as more loci were included. Maximum likelihood (ML) trees were reconstructed under a codon-partitioned scheme, with results shown in Supplementary Figs. S16–20 (500 loci) and S21–25 (700 loci). Within Elasmobranchii, topologies remained consistent with those from the 300-locus datasets. Notably, only the dataset filtered by absRatioLen recovered Selachii as monophyletic (Supplementary Fig. S26), likely due to this metric’s emphasis on balancing ingroup and outgroup branch lengths. In contrast, relationships within Batoidea remained highly discordant, although most trees supported the Batoidea-1 topology. Overall, increasing the number of loci did not affect the stability of Elasmobranch relationships. All filtering strategies recovered consistent topologies, with only the absRatioLen-based dataset resolving Selachii as monophyletic. The 300-locus threshold appears to provide an optimal balance: it captures the highest proportion of unique loci and corresponds to the point before the signal distribution turning point (Fig. S7).

### Topology-Based Filtering Methods

Topology-based filtering methods complement metric-based approaches in phylogenomic studies by discover genes with weak phylogenetic signals or those exhibiting extreme likelihood values, thereby refining species tree inference (Hughes et al., 2023; Li et al., 2021; Shen et al., 2017; Smith et al., 2020). These methods are particularly valuable for resolving conflicting relationships that arise when alternative evolutionary models or inference methods yield incongruent topologies at contentious nodes. By constraining phylogenetic hypotheses to predefined topologies, researchers can compute gene-specific likelihood scores, statistically compare support for competing hypotheses, and evaluate the impact of excluding outlier loci on tree reconstruction (Li et al., 2021; Shen et al., 2017; Smith et al., 2020).

A widely adopted approach, the Gene-wise Log-likelihood Score (GLS), quantifies individual gene support for alternative topologies, identifying loci that disproportionately influence species relationships (Shen et al., 2017). Studies have demonstrated that even one or two outlier genes with large likelihood differences (ΔGLS) can dramatically alter inferred phylogenies (Brown & Thomson, 2017; Walker et al., 2018). Similarly, methods like Gene Genealogy Interrogation (GGI) account for gene tree estimation error by constraining analyses to a predefined tree space, often encompassing all possible topologies for a focal lineage, to test specific evolutionary hypotheses (Arcila et al., 2017).

In this study, we applied GLS to investigate unresolved relationships within Elasmobranchii, focusing on three major superorders: Galeomorphii, Squalomorphii, and Batoidea. Based on literature-derived hypotheses, we tested three alternative topologies (Fig. 1): (1) Elasmobranchii-1: Selachii (sharks) as sister to Batoidea (rays/skates); (2) Elasmobranchii-2: Squalomorphii as sister to Batoidea + Galeomorphii; (3) Elasmobranchii-3: Galeomorphii as sister to Batoidea + Squalomorphii. Initial analyses of unfiltered datasets recovered Elasmobranchii-1 under coalescent methods and Elasmobranchii-2 under concatenation (Fig. 4). Using these topologies as constraints, we calculated ΔGLS values across loci in the NT dataset-1 to assess gene-level support. Outlier loci were defined as those with ΔGLS values beyond ±20, a threshold determined empirically from the ΔGLS distribution (Fig. 5c). Exclusion of nine outlier loci and subsequent reanalysis produced topologies identical to those from unfiltered data, suggesting minimal influence of these loci on overall resolution. This aligns with findings by Hughes et al. (2023), who found little difference in gene-tree likelihood scores between the two hypthetical topologies and showed that removing outliers had minimal impact.

A key limitation of topology-based methods lies in their reliance on predefined phylogenetic hypotheses. Additionally, ΔGLS filtering may fail to resolve cases where likelihood differences are symmetrically distributed across loci.

### Resolving Chondrichthyan Phylogenetic Relationships

Our phylogenomic analyses provide crucial insights into three long-standing controversies in chondrichthyan evolution: (1) the monophyly of Squalomorphii, (2) the reciprocal monophyly of sharks (Selachii) and rays (Batoidea), and (3) relationships within Batoidea. The combined molecular and morphological evidence strongly supports Hexanchiformes as the basal lineage within Squalomorphii, consistent with previous mitochondrial (Naylor et al., 2012) and targeted gene analyses (Straube et al., 2015). This placement was particularly robust in our absRatioLen-filtered analyses.

The monophyly of Squalomorphii was consistently recovered from earlier mitochondrial and limited nuclear gene studies, but was rejected in phylogenomic dataset, even under traditional metric-based filtering method. It placed Hexanchiformes as paraphyletic to the rest of Squalomorphii. Our novel filtering approach, absRatioLen, adjusted this placement and recovered back as monophyly. However, a notable exception arose in the coalesecnt analysis with absRatioLen, which weakly supported Pristiophoriformes as sister to Squaliformes (posterior probability = 0.69; Supplementary Fig. S12). This unexpected result may reflect limited taxon sampling within Pristiophoriformes. Additionally, analyses including Echinorhiniformes with the 88-locus dataset positioned it as sister to a clade of Pristiophoriformes and Squatiniformes, with Squaliformes as their closest sister group, while Hexanchiformes remained paraphyletic (Supplementary Figs. S14–S15). This unusual position is likely caused by insufficient dataset which is unable to conduct filtering step.

Regarding the shark-ray dichotomy, our results conclusively reject the morphological hypnosqualean hypothesis (Carvalho, 1996; Shirai, 1996) that nested Batoidea within Squalomorphii. Instead, we recovered Selachii and Batoidea as reciprocally monophyletic groups, aligning with recent integrative analyses combining NADH2 sequences with pelvic morphology (da Silva & Vaz, 2023) and supporting their status as distinct evolutionary lineages since at least the Triassic (Maisey, 2012). This finding was particularly consistent in absRatioLen-filtered datasets, which overcame the topological instability observed in unfiltered analyses and traditional metric approaches.

Within Batoidea, our results reveal significant conflict between molecular and morphological evolutionary hypotheses. While recent morphological analyses position Torpediniformes as sister to Myliobatiformes at the distal tips of the batoid tree (Villalobos-Segura et al., 2022), our molecular data consistently support Rajiformes as the basal group with Torpediniformes sister to Rhinopristiformes + Myliobatiformes—a topology also recovered in most nuclear and mitochondrial studies (Naylor, 2025; Stein et al., 2018). These discrepancies likely reflect the challenges of establishing clear synapomorphies in batoid morphology and underscore the value of phylogenomic approaches for clarifying their evolutionary history.

The success of our absRatioLen filtering method in resolving these longstanding conflicts demonstrates the critical importance of branch length metrics for detecting systematic errors in deep phylogenetic reconstruction. Unlike traditional approaches or GC-content based filtering (absDiffGC), absRatioLen effectively captured lineage-specific evolutionary patterns while minimizing artifacts like long-branch attraction. This was particularly evident in its ability to stabilize the position of problematic groups like Hexanchiformes and produce topologies congruent with the growing consensus from multiple molecular datasets.

### Resolving Deep Vertebrate Phylogenetic Conflicts beyond Chondrichthyans

Our absRatioLen filtering method demonstrates broad applicability beyond chondrichthyans by successfully resolving one of vertebrate phylogeny’s most prominent conflicts, the relationships among lungfish, coelacanths, and tetrapods. This triad represents a classic case of phylogenomic incongruence, with competing topologies recurrently proposed despite extensive genomic sampling (Irisarri et al., 2017; Takezaki & Nishihara, 2017). Notably, our filtered dataset consistently recovered lungfish as the sister group to tetrapods (Supplementary Fig. S33F), even when using outgroup configurations that previously produced conflicting results (Takezaki & Nishihara, 2017). This robust resolution underscores a critical advantage of branch-length based filtering: its ability to overcome systematic errors induced by outgroup selection and evolutionary rate heterogeneity.

The success of absRatioLen in this challenging case study highlights its potential as a general solution for phylogenomic conflicts across taxonomic scales. Similar incongruences plague diverse systems, from deep divergences in Ostariophysan fishes (Chakrabarty et al., 2017) to recent radiations like flatfish origins within Carangaria (Duarte-Ribeiro et al., 2024; Lü et al., 2021). Our results suggest these conflicts may often stem from localized signal erosion rather than true biological complexity, a hypothesis supported by absRatioLen’s capacity to recover consistent topologies by selectively retaining loci with balanced evolutionary rates. This approach proves particularly valuable when traditional methods yield conflicting results, offering a principled framework for data curation prior to phylogenetic inference.

## Conclusions

Our study reveals fundamental limitations in traditional phylogenomic approaches for resolving deep evolutionary relationships in Chondrichthyes. While mitochondrial and limited nuclear datasets have provided preliminary insights, they consistently fail to resolve key nodes with strong support. We demonstrate that conventional filtering methods, whether metric-based or topology-driven, share substantial parameter overlap and consequently show limited efficacy for addressing deep phylogenetic conflicts. The novel branch ratio method (absRatioLen) overcomes these limitations by focusing on localized evolutionary signals rather than genome-wide patterns. This approach consistently recovered robust, consensus topologies across analytical frameworks, including the critical case of Hexanchiformes placement. By targeting clade-specific heterogeneity rather than global gene properties, absRatioLen provides a powerful solution to persistent systematic discordance in chondrichthyan phylogenomics. Our findings establish branch length ratio filtering as a transformative approach for resolving deep evolutionary relationships. This method not only clarifies long-standing controversies in chondrichthyan systematics but also offers a general framework for investigating contentious nodes across the tree of life. The success of this localized signal approach opens new avenues for reconciling molecular phylogenetics with morphological and ecological diversification patterns in ancient vertebrate lineages.

## Notes

### Competing Interest Statement

The authors have declared no competing interest.

