## Supplementary Figure for "A novel data filtering method resolves the controversy in the phylogeny of the Chondrichthyes"

This file contains Supplementary Figures S1-S33. See Excel file for Supplementary Table S1-7.

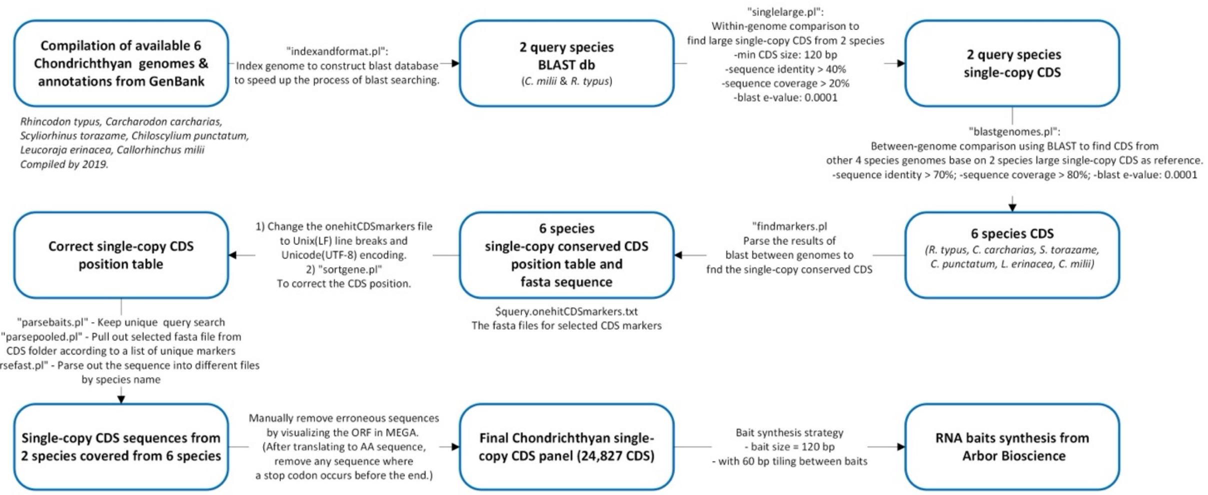


Fig. S1. Flowchart illustrating the Chondrichthyan bait design process. Six publicly available Chondrichthyan genomes, compiled by 2019, were analyzed to identify single-copy coding sequences (CDSs) for bait development. 24,827 CDSs were selected and synthesized into 120 bp baits by Arbor Bioscience.

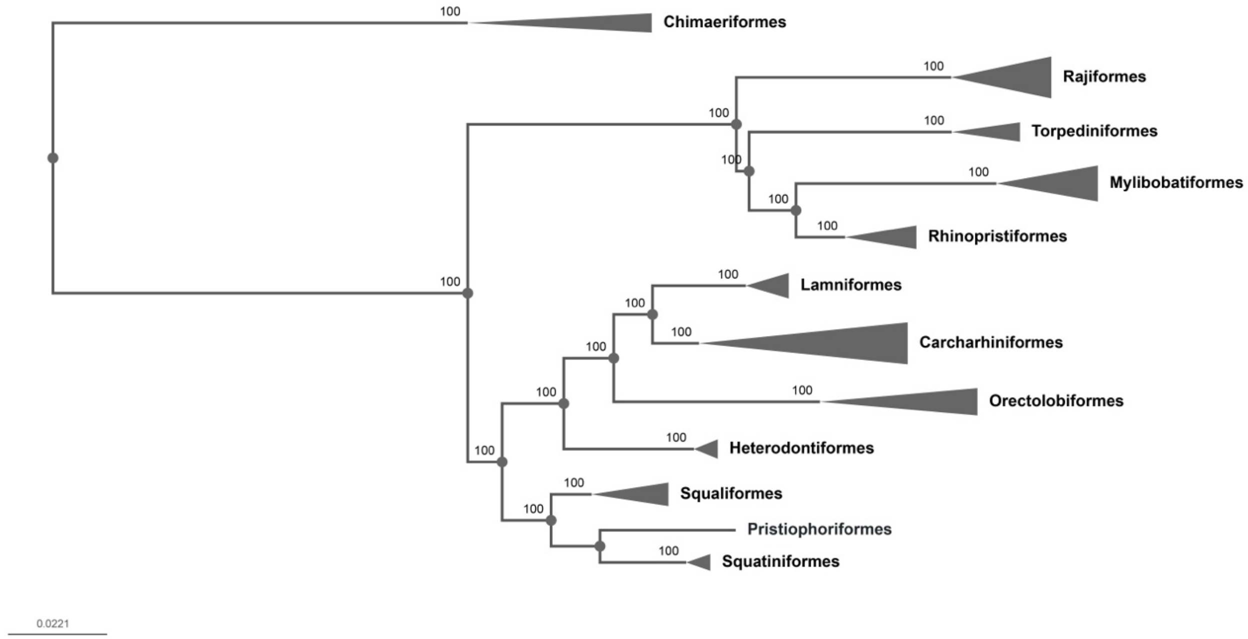


Fig. S2. Maximum likelihood phylogram inferred using IQ-TREE from the NT-Hex data matrix under the partition-by-codon scheme, visualized in phylo.io.

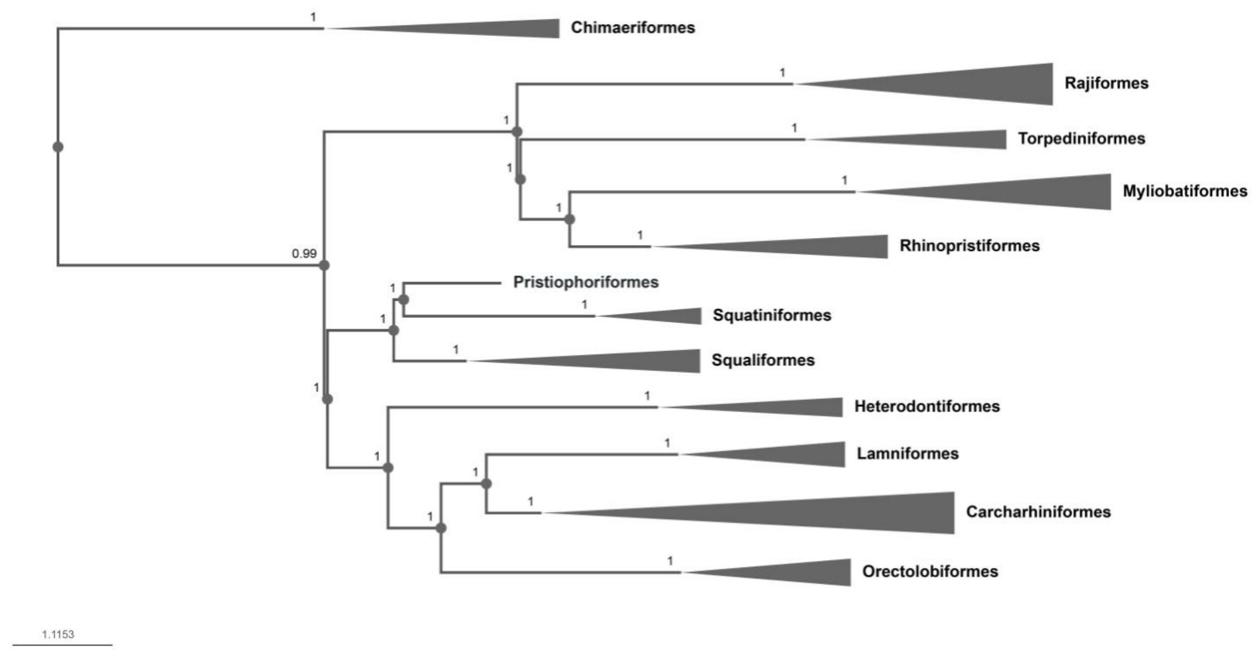


Fig. S3. Multispecies Coalescent phylogram inferred using ASTRAL from the NT-Hex data matrix, visualized in phylo.io.

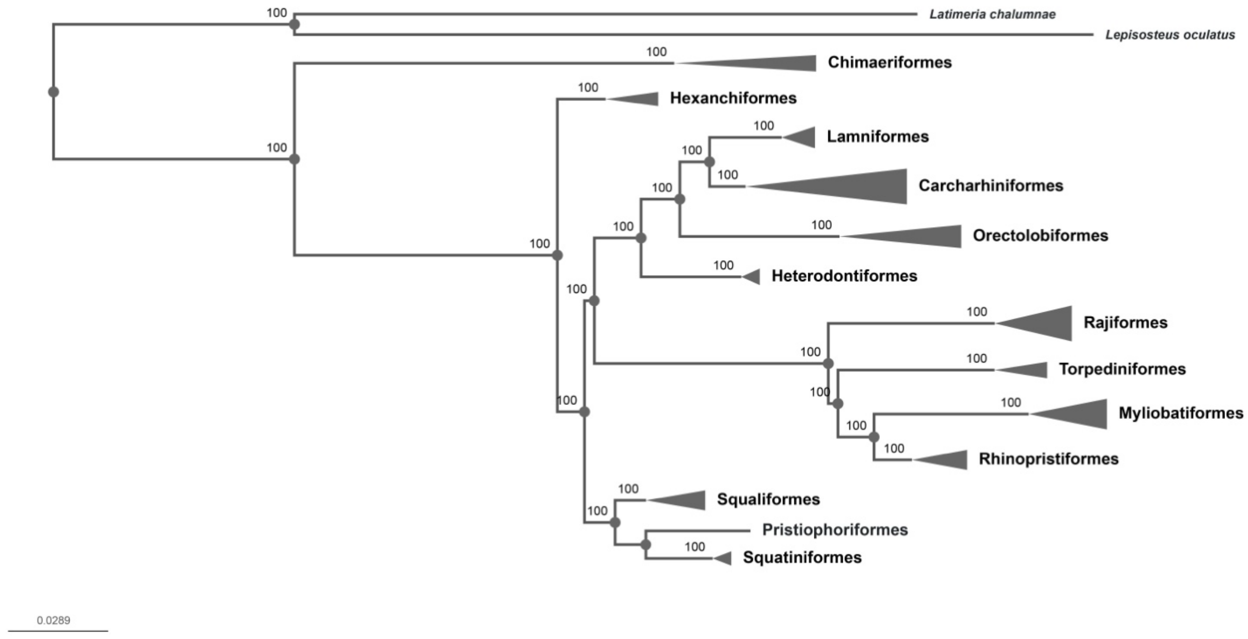


Fig. S4. Maximum likelihood phylogram inferred using IQ-TREE from the NT-Out data matrix under the partition-by-codon scheme, visualized in phylo.io.

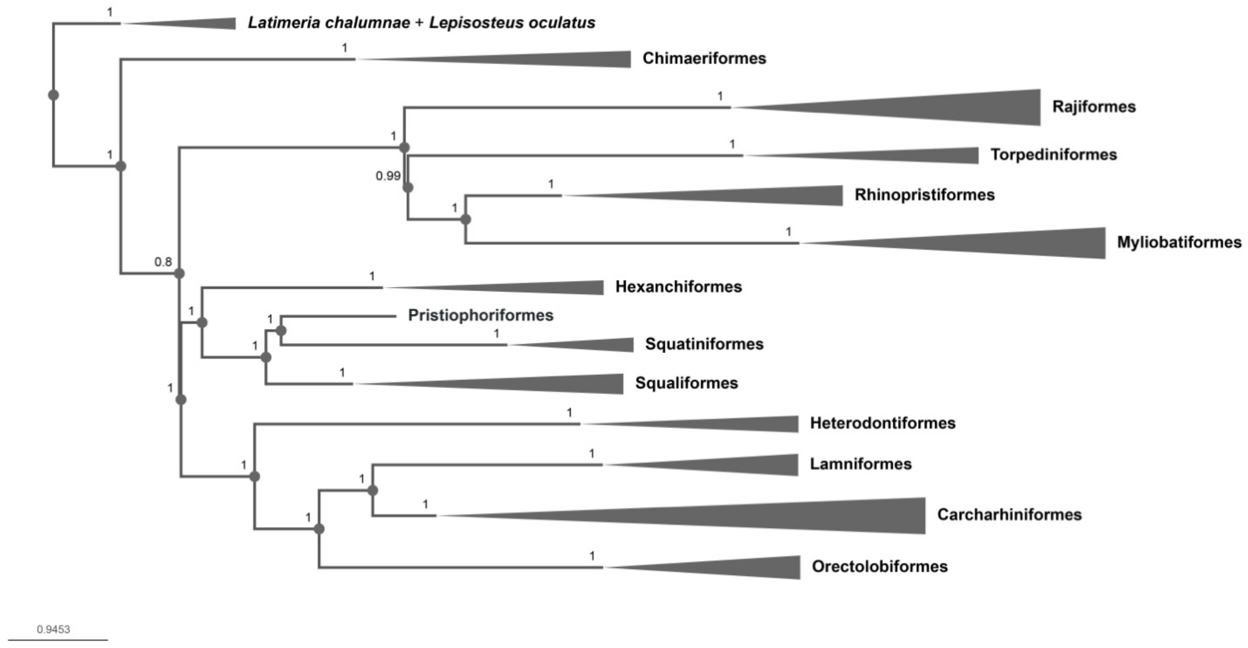


Fig. S5. Multispecies Coalescent phylogram inferred using ASTRAL from the NT-Out data matrix, visualized in phylo.io.

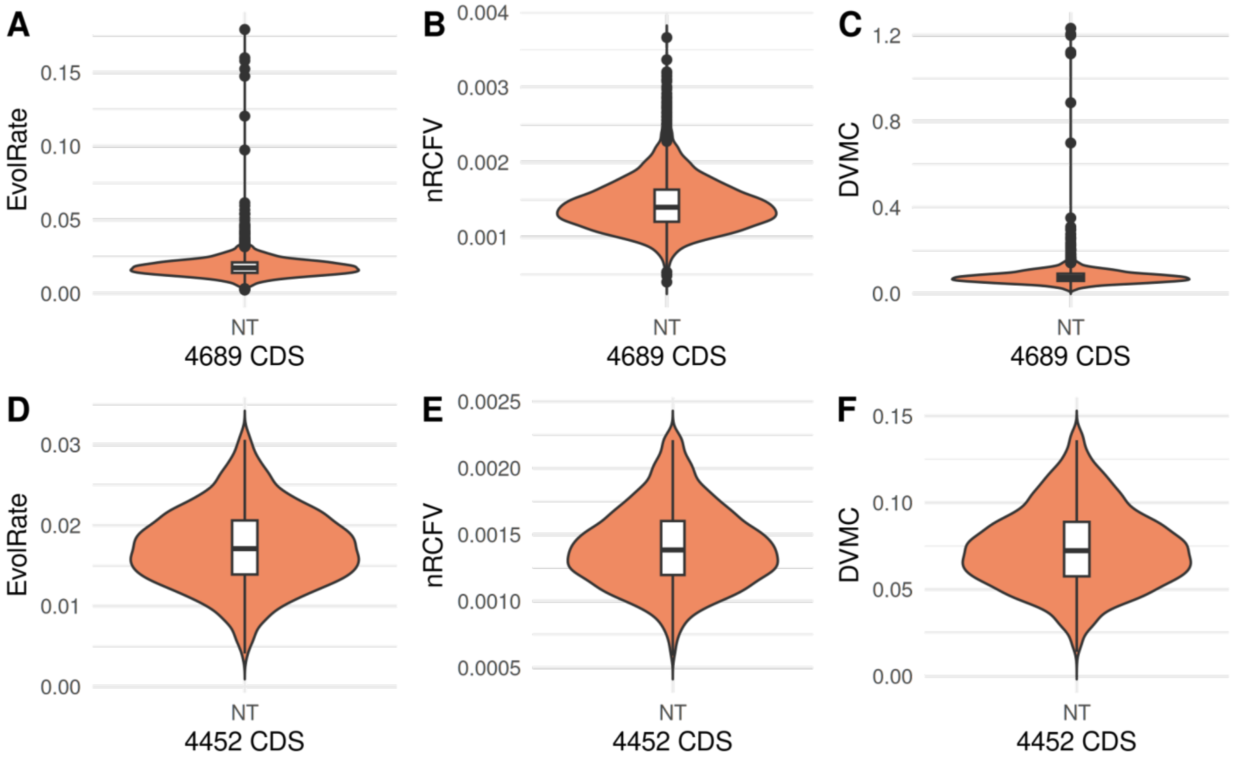


Fig. S6. Violin plots, nested with box plots, showing the distribution of three phylogenetic signals before and after filtering. The pre-filtered dataset includes the low PIS discaded matrix (4,689 CDS), while the post-filtered dataset represents the phylogenetic signal outliers removed matrix (4,452 CDS). The bold line within each box plot indicates the median, and solid black dots represent outliers that exceed 1.5 times the interquartile range. It is important to note that points falling outside the 1.5× interquartile range in filtered dataset are not necessarily outliers in the unfiltered dataset. To avoid misinterpretation, such points are not displayed as black dots.

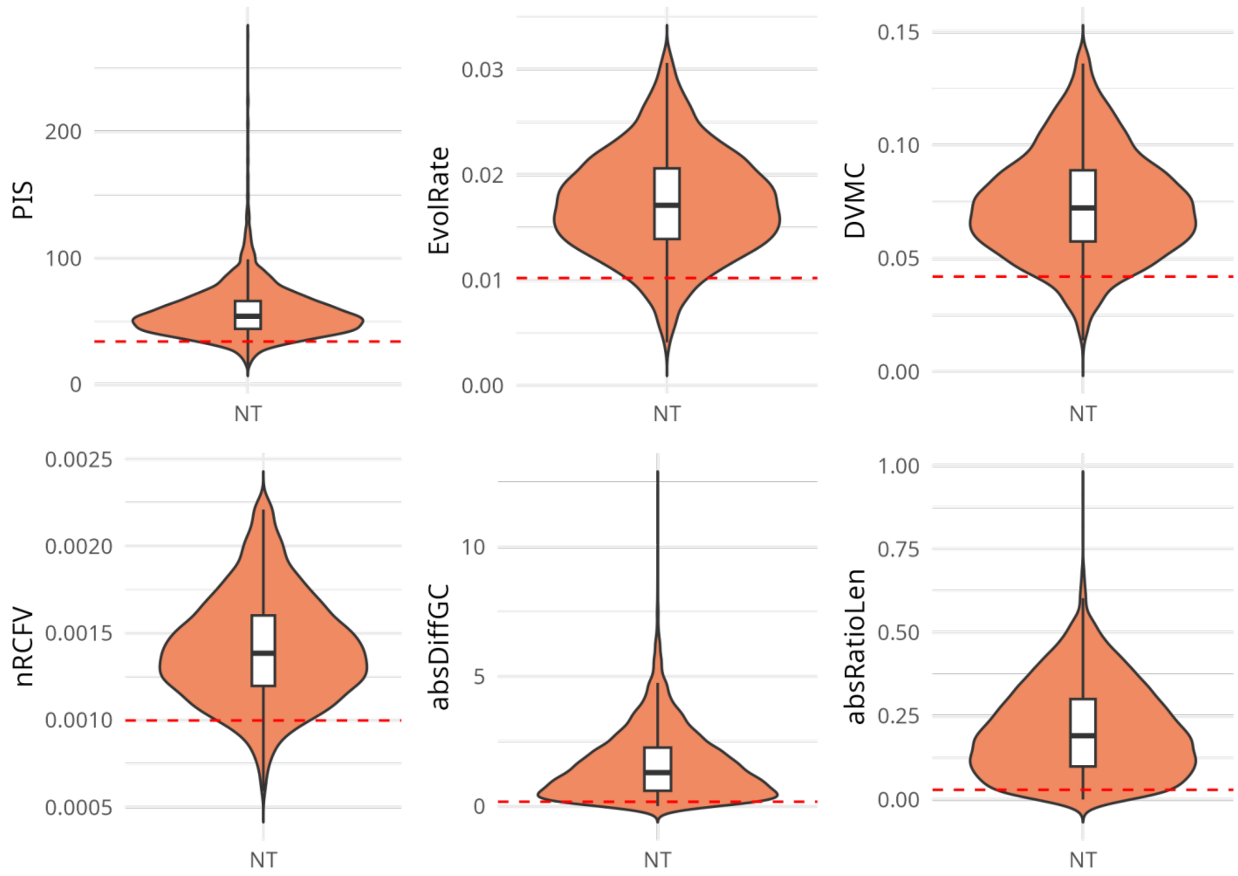


Fig. S7. Violin plots, nested with box plots, illustrating the distribution of phylogenetic signals in the NT dataset (4,452 CDS) from four traditional and two novel phylogenetic signals. The bold line within each box represents the median. “Outlier” points are not displayed, as they are not outliers when compared to the standard in Supplementary Figure S2. This intentional omission prevents potential misinterpretation of “outliers” points. The red dashed line indicates the 300$^{th}$ ranked data point.

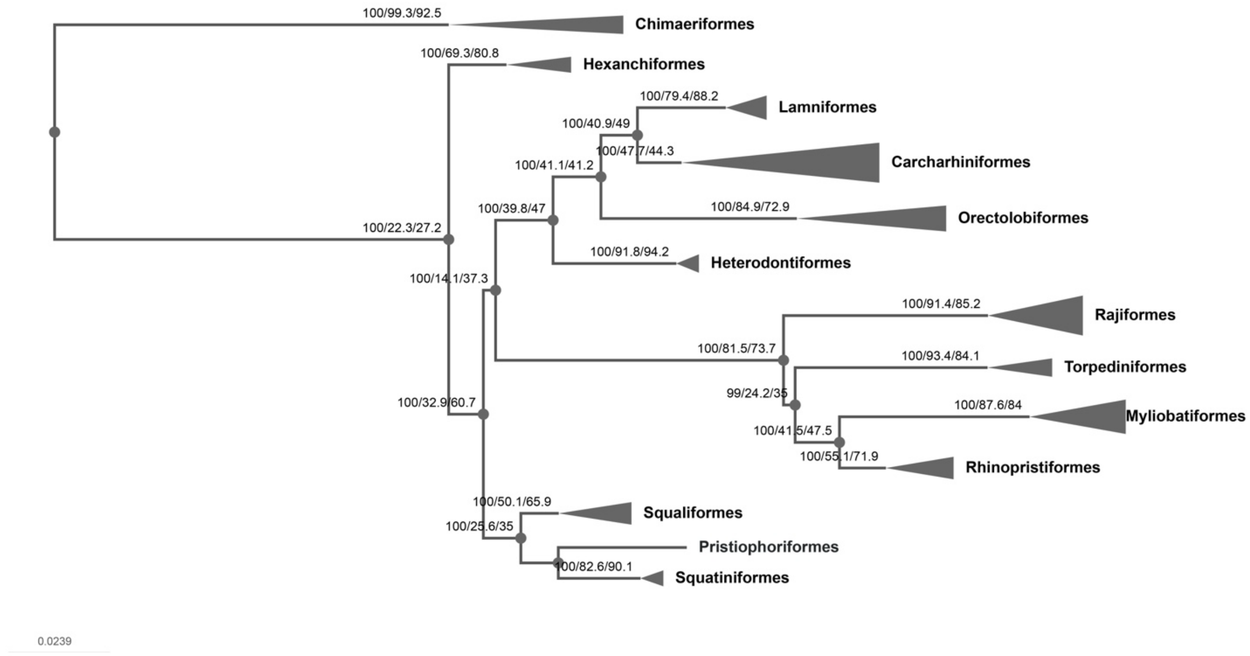


Fig. S8. Maximum likelihood phylogram inferred using IQ-TREE from the clean NT data matrix under the partition-by-codon scheme, visualized in phylo.io (Robinson et al., 2016). Branch labels indicate bootstrap support values, gene concordance factors (gCF), and site concordance factors (sCF). Monophyletic clades are represented as wedge-shaped clusters, except for *Pristiophorus cirratus*, which only has a single sample.

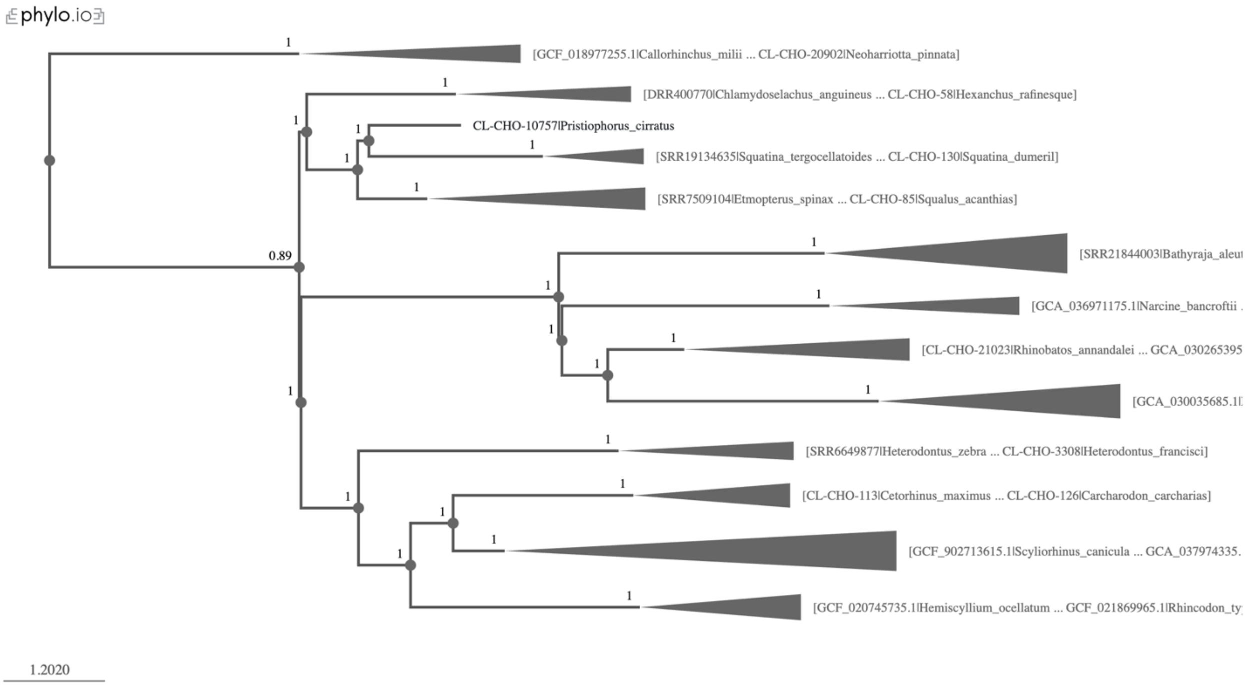


Fig. S9. Multispecies Coalescent phylogram inferred using ASTRAL from the clean NT data matrix, visualized in phylo.io.

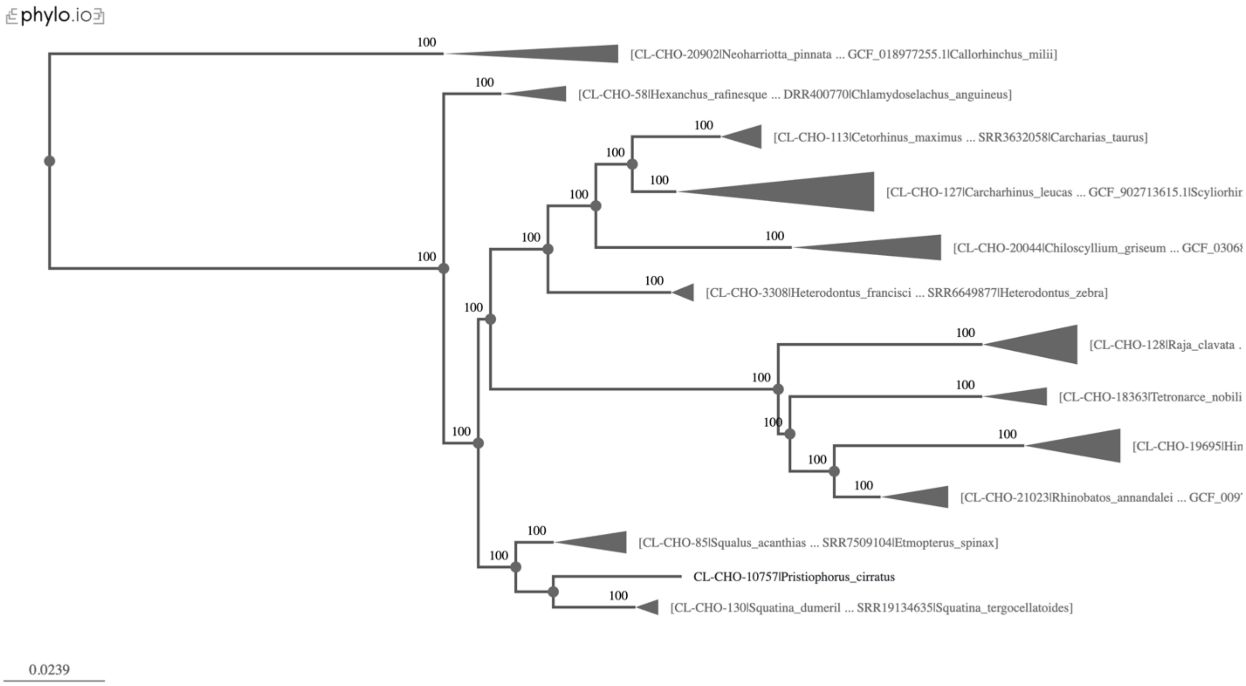


Fig. S10. Maximum likelihood phylogram inferred using IQ-TREE from the NT without GLS outliers data matrix under the partition-by-codon scheme, visualized in phylo.io

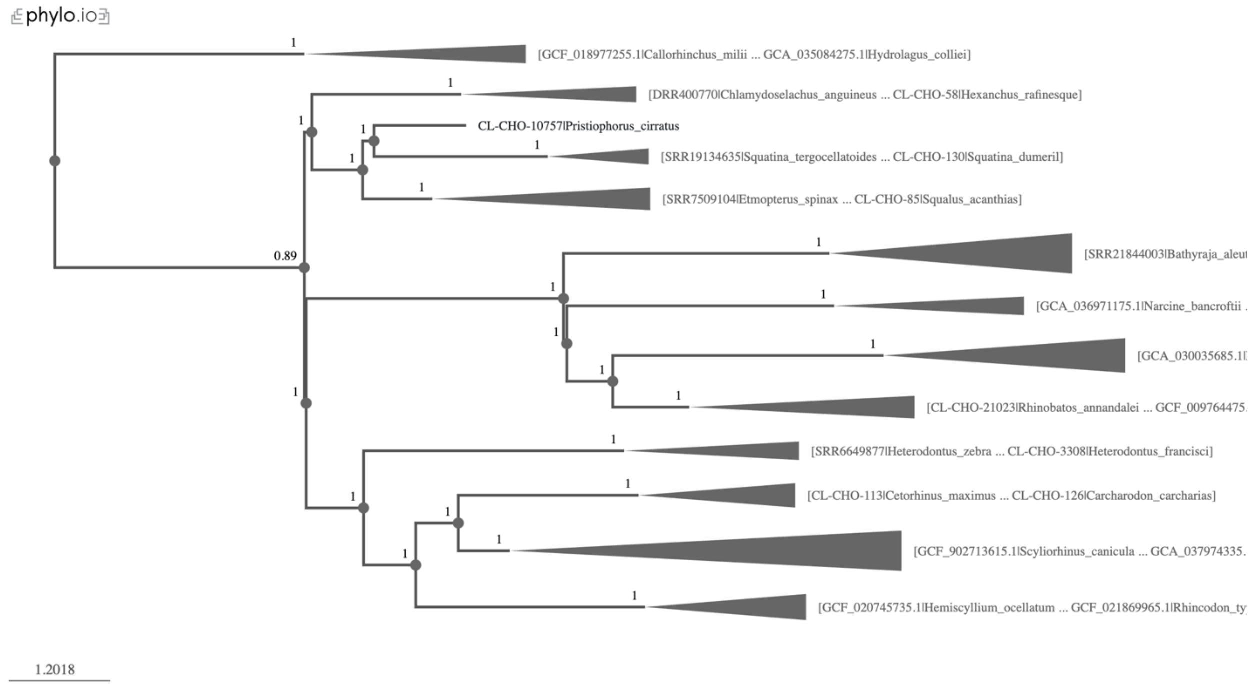


Fig. S11. Multispecies Coalescent phylogram inferred using ASTRAL from the NT without GLS outliers data matrix, visualized in phylo.io.

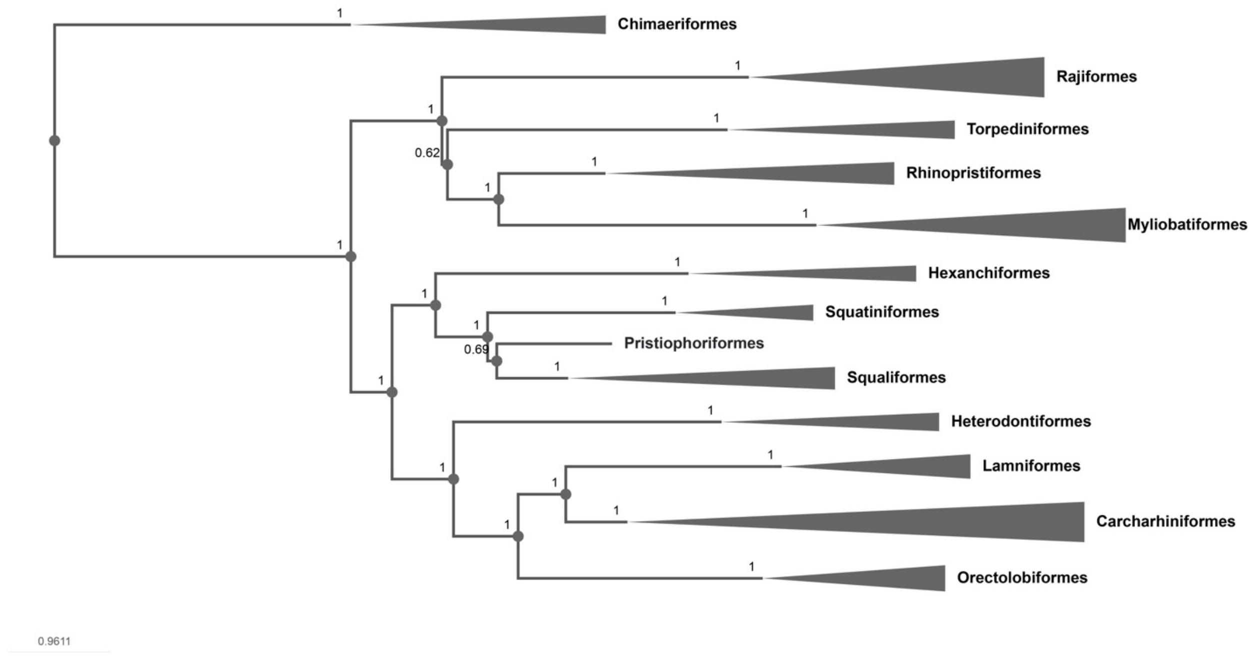


Fig. S12. Multispecies Coalescent phylogram inferred using ASTRAL from the NT-300absRatioLen data matrix, visualized in phylo.io.

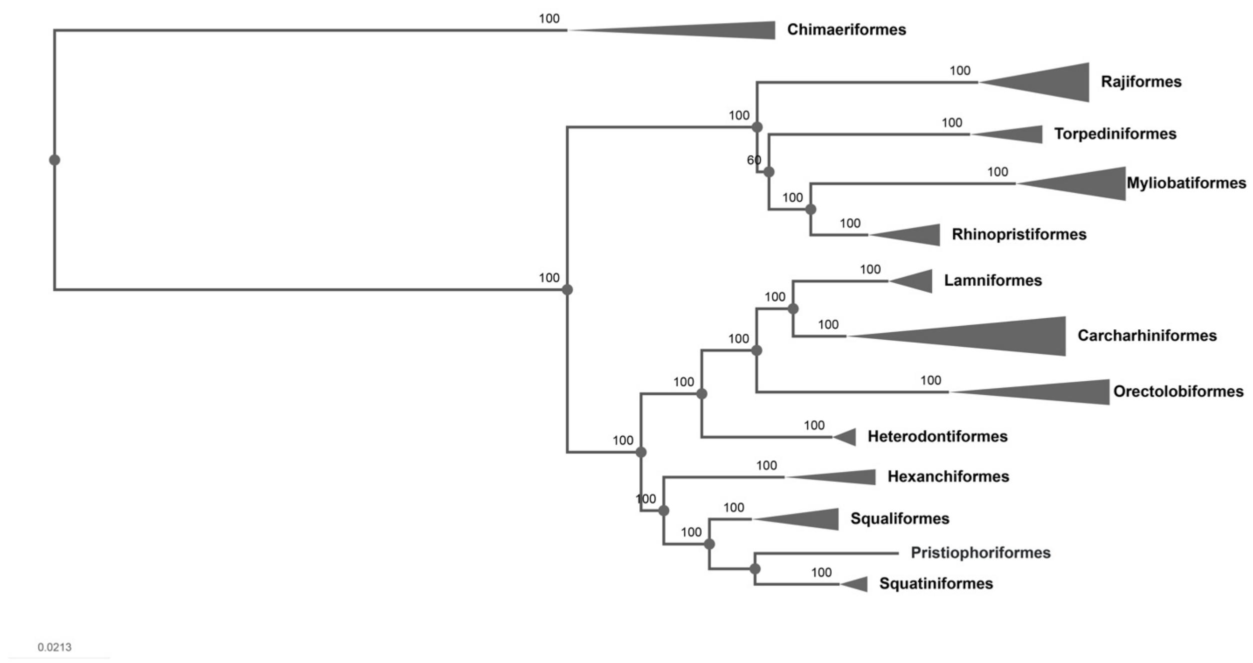


Fig. S13. Maximum likelihood phylogram inferred using IQ-TREE from the NT-300absRatioLen data matrix under the partition-by-codon scheme, visualized in phylo.io.

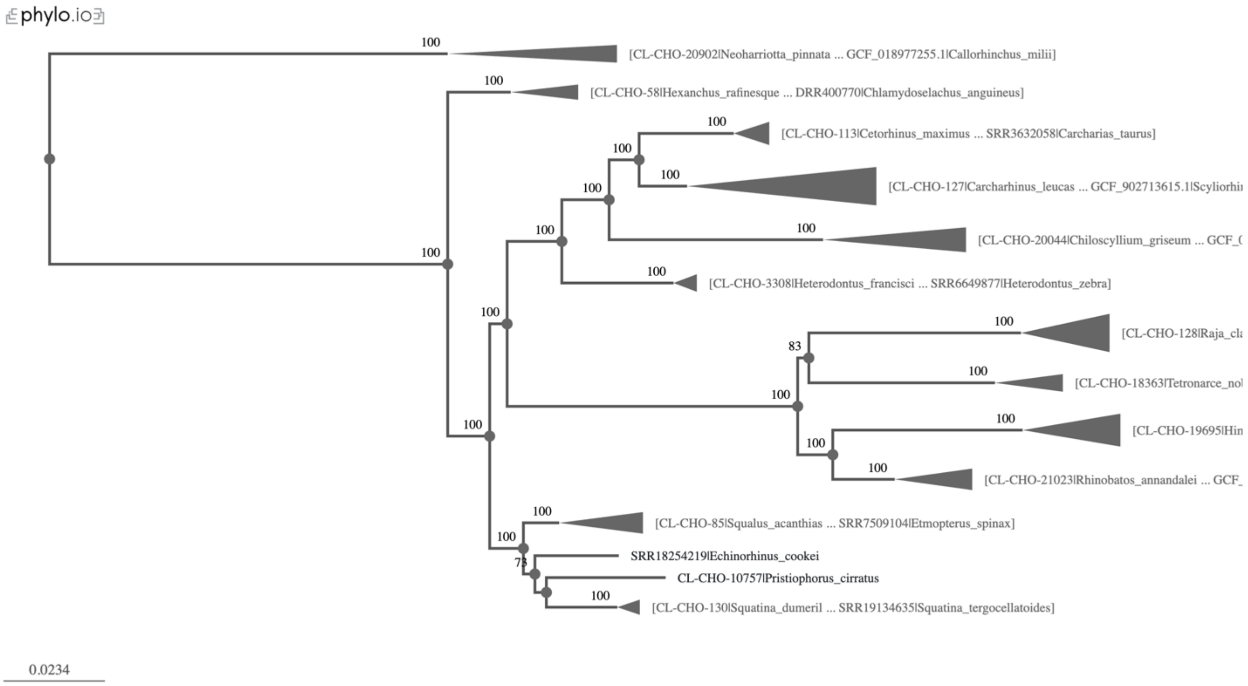


Fig. S14. Maximum likelihood phylogram inferred using IQ-TREE from the NT-ech data matrix (88 loci) under the partition-by-codon scheme, visualized in phylo.io.

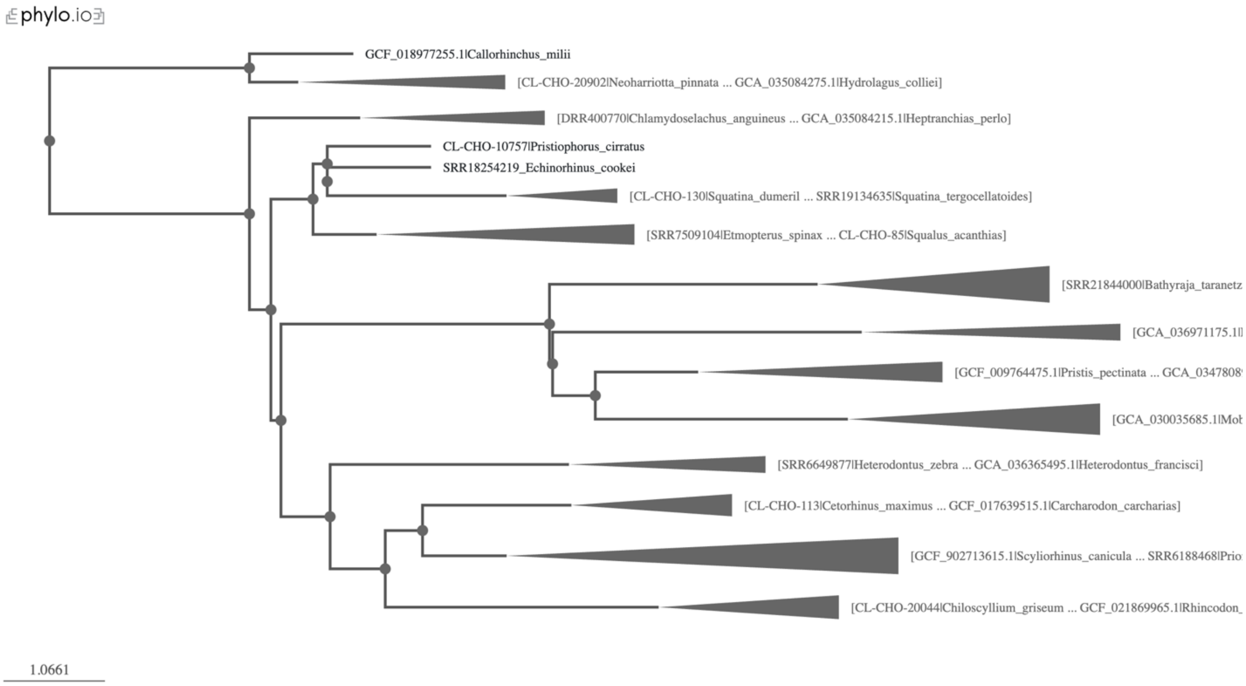


Fig. S15. Multispecies Coalescent phylogram inferred using ASTRAL from the NT-ech data matrix (88 loci), visualized in phylo.io.

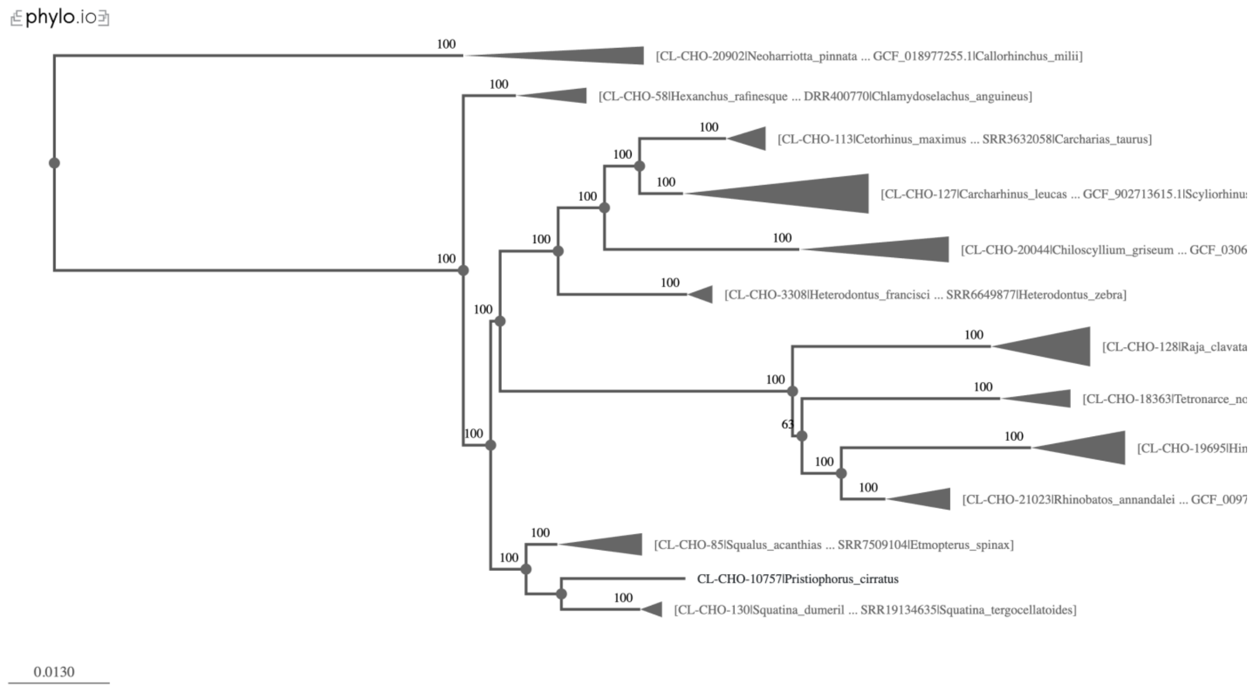


Fig. S16. Maximum likelihood phylogram inferred using IQ-TREE from the NT-500evoRate data matrix under the partition-by-codon scheme, visualized in phylo.io.

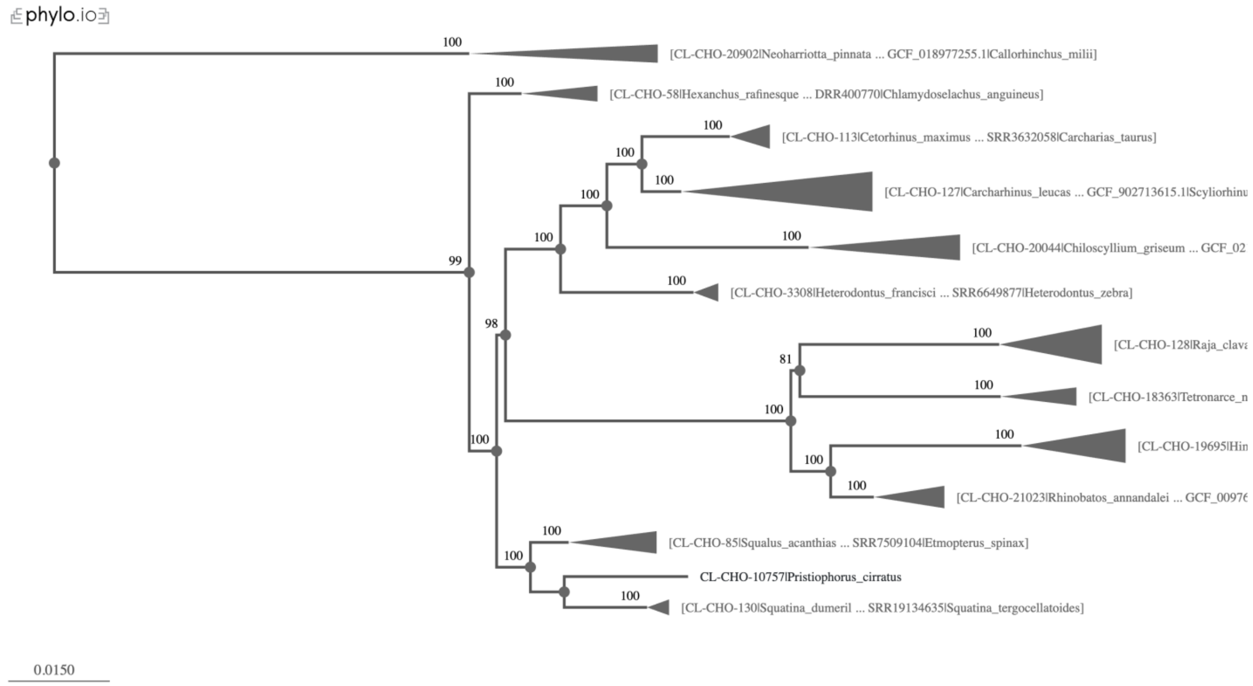


Fig. S17. Maximum likelihood phylogram inferred using IQ-TREE from the NT-500nrcfv data matrix under the partition-by-codon scheme, visualized in phylo.io.

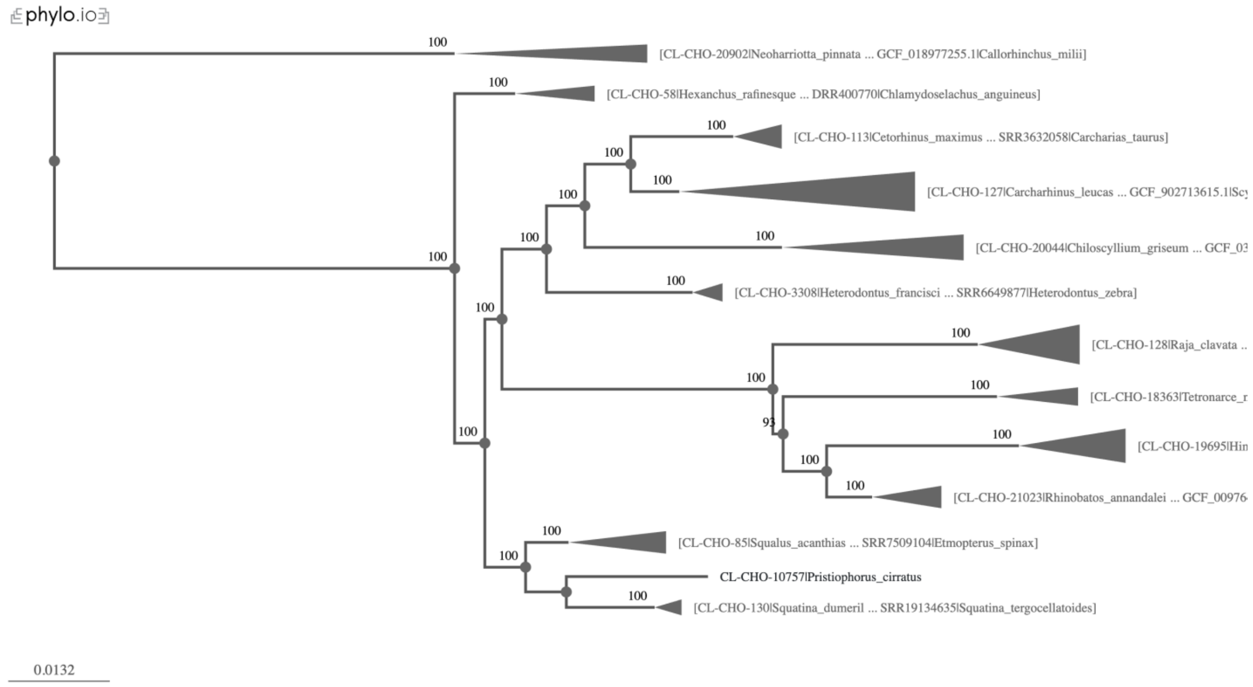


Fig. S18. Maximum likelihood phylogram inferred using IQ-TREE from the NT-500dvmc data matrix under the partition-by-codon scheme, visualized in phylo.io.

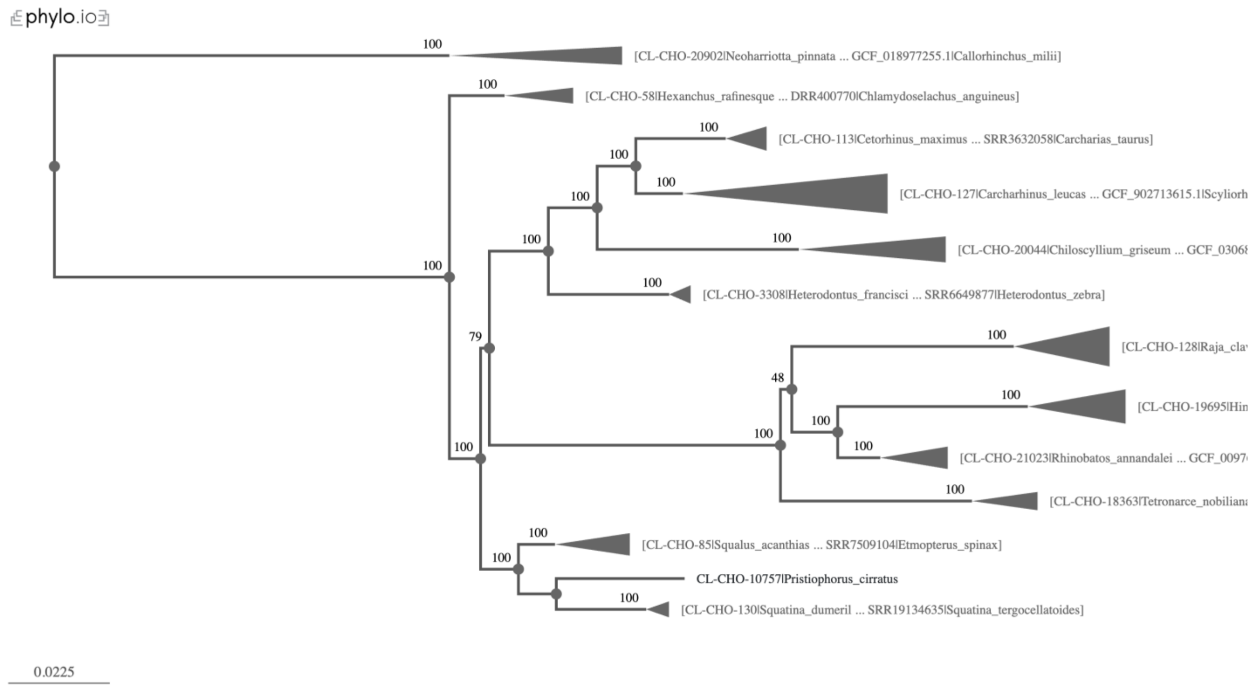


Fig. S19. Maximum likelihood phylogram inferred using IQ-TREE from the NT-500absDiffGC data matrix under the partition-by-codon scheme, visualized in phylo.io.

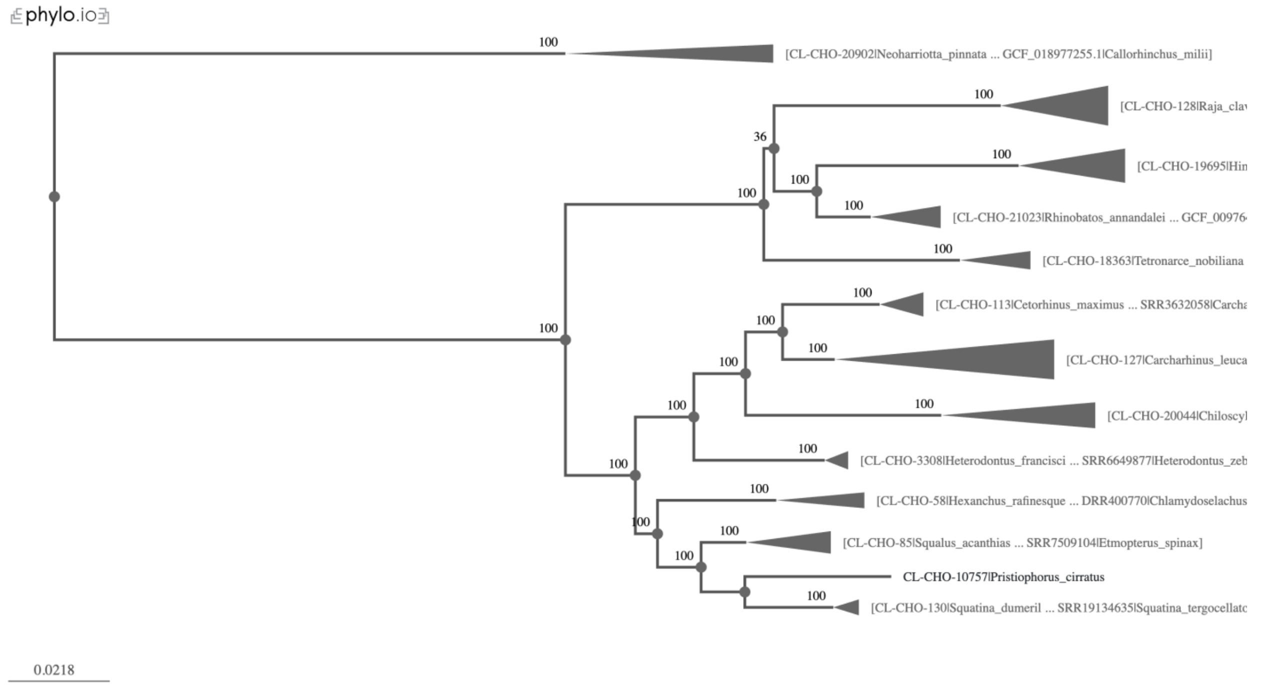


Fig. S20. Maximum likelihood phylogram inferred using IQ-TREE from the NT-500absRatioLen data matrix under the partition-by-codon scheme, visualized in phylo.io.

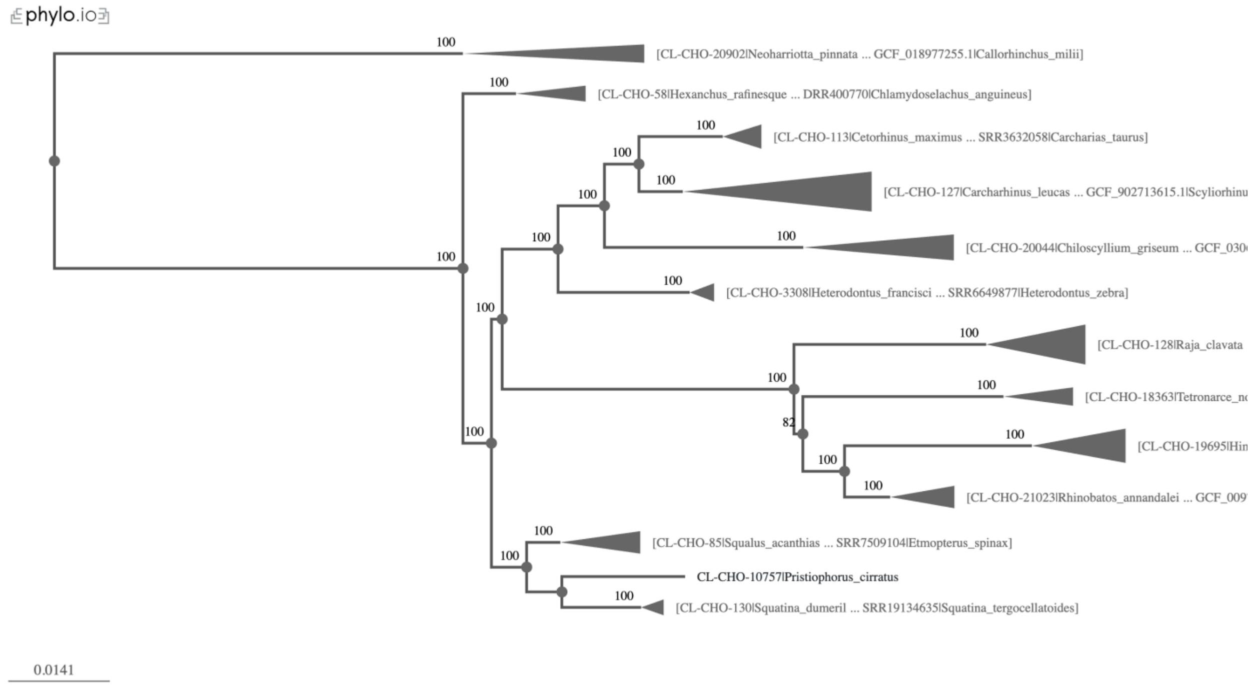


Fig. S21. Maximum likelihood phylogram inferred using IQ-TREE from the NT-700evoRate data matrix under the partition-by-codon scheme, visualized in phylo.io.

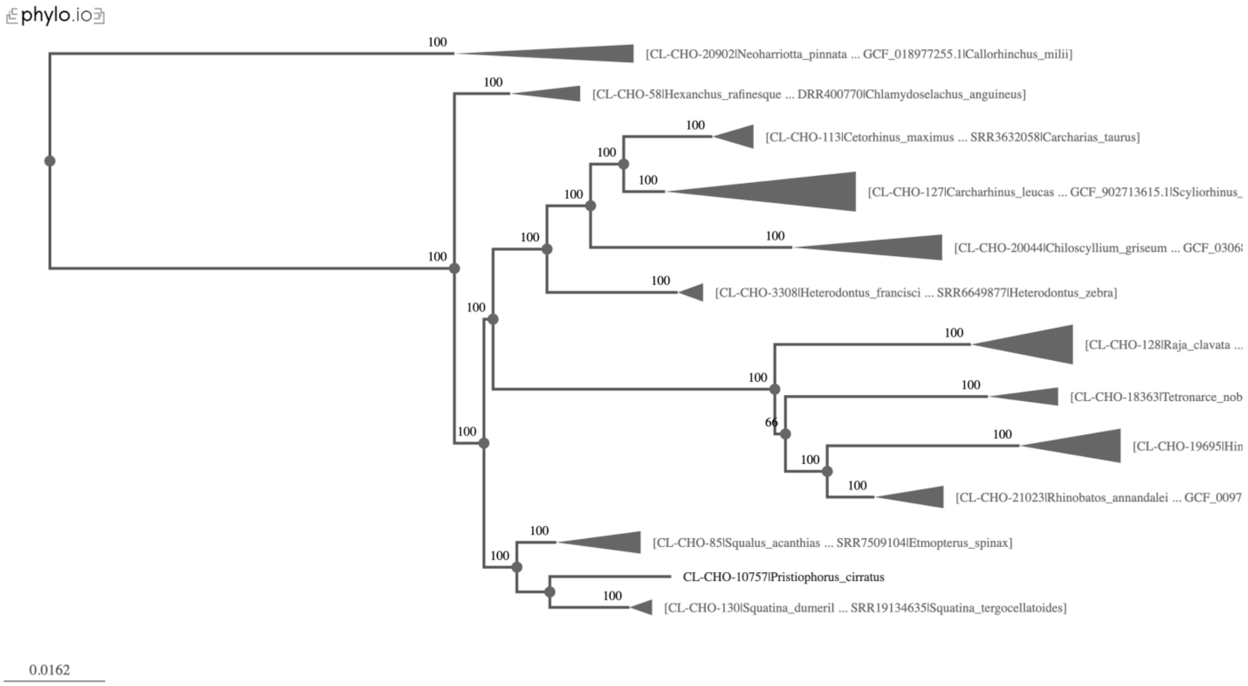


Fig. S22. Maximum likelihood phylogram inferred using IQ-TREE from the NT-700nrcfv data matrix under the partition-by-codon scheme, visualized in phylo.io.

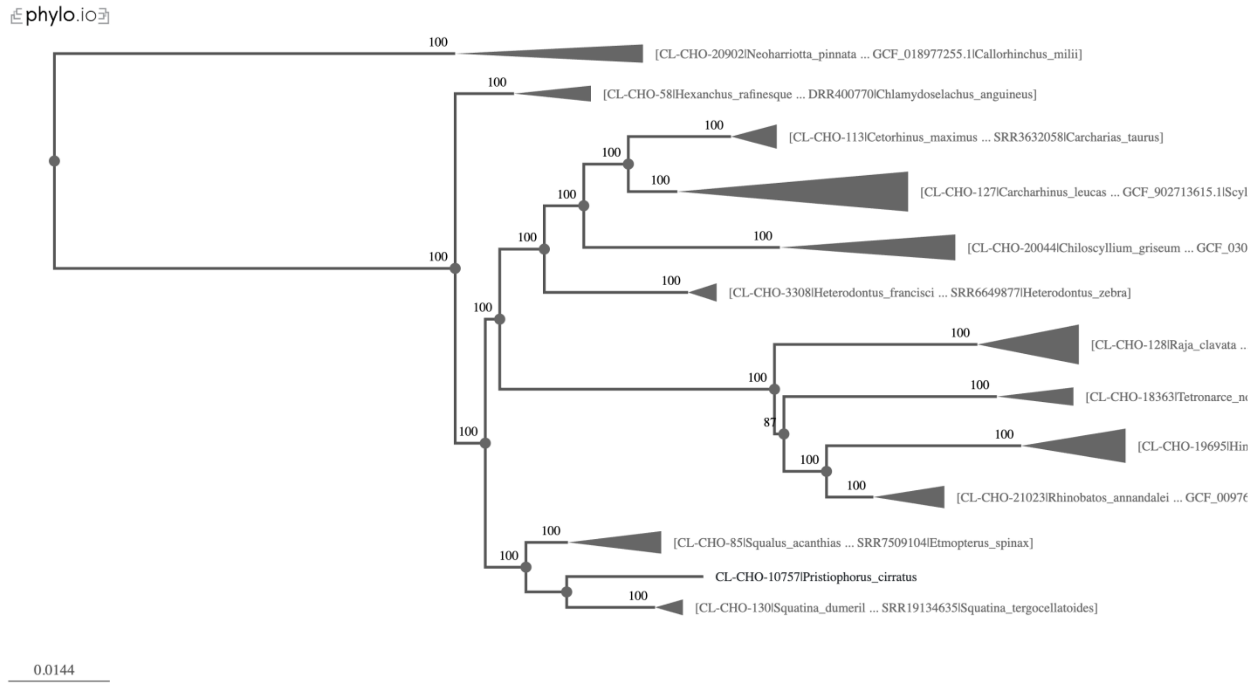


Fig. S23. Maximum likelihood phylogram inferred using IQ-TREE from the NT-700dvmc data matrix under the partition-by-codon scheme, visualized in phylo.io.

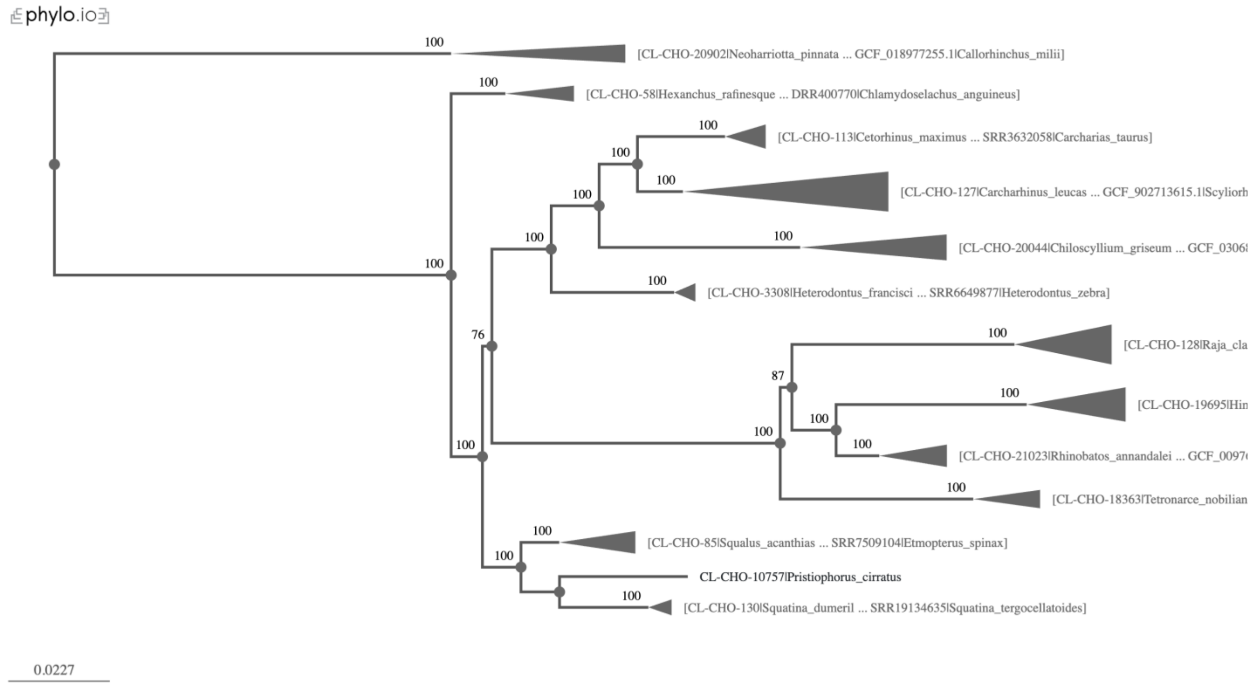


Fig. S24. Maximum likelihood phylogram inferred using IQ-TREE from the NT-700absDiffGC data matrix under the partition-by-codon scheme, visualized in phylo.io.

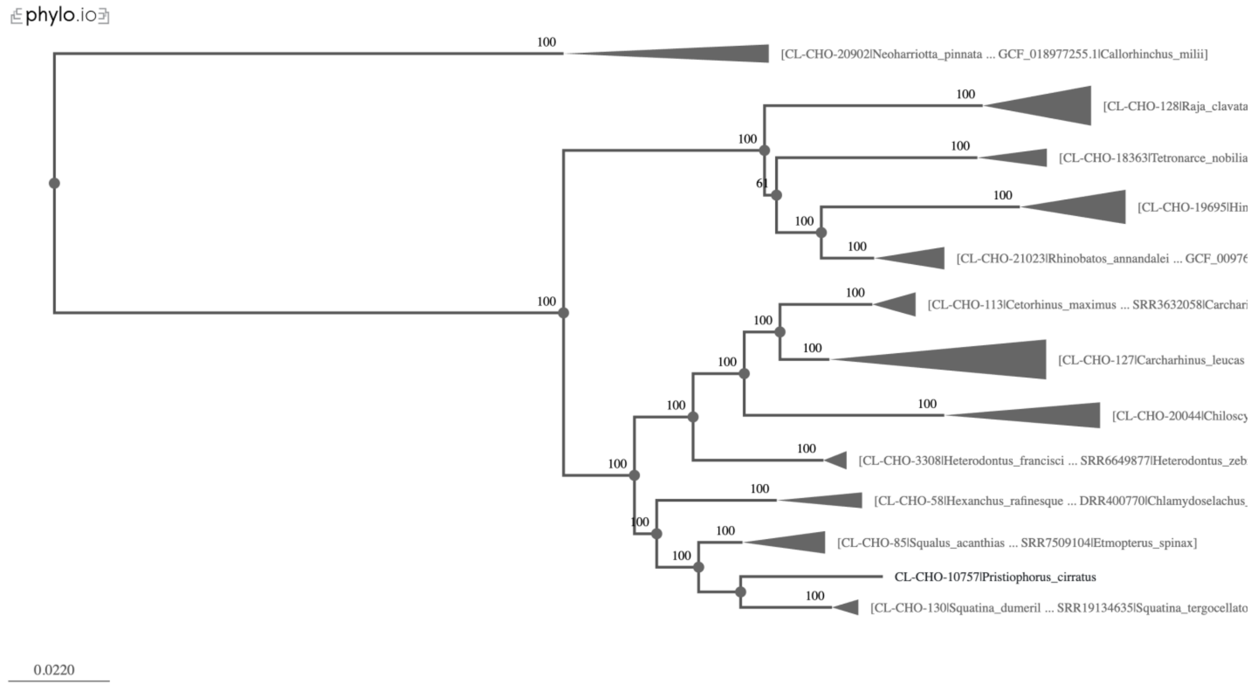


Fig. S25. Maximum likelihood phylogram inferred using IQ-TREE from the NT-700absRatioLen data matrix under the partition-by-codon scheme, visualized in phylo.io.

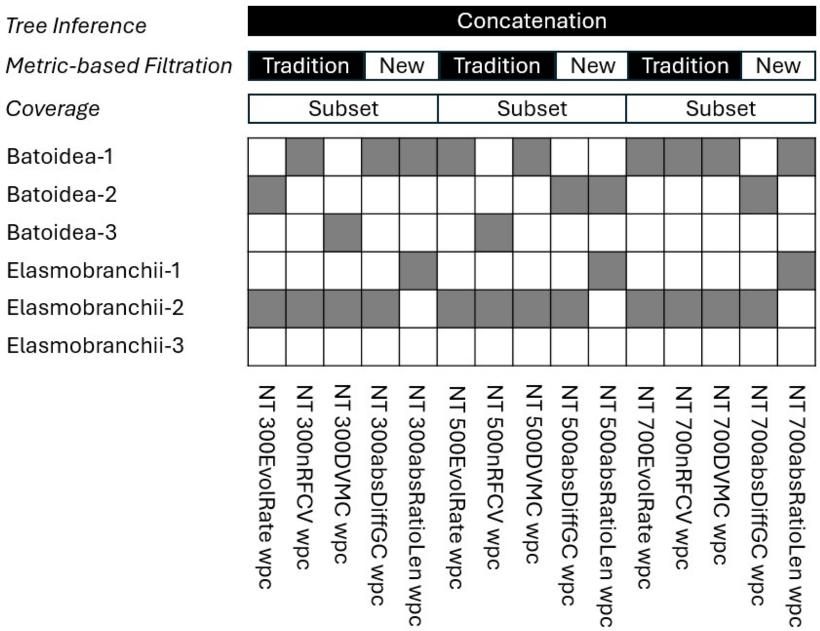


Fig. S26. Summary of topological support for controversial Elasmobranchii and Batoidea relationships across 15 phylogenetic reconstructions from subset dataset filtering by 3 traditional metric and 2 novel metric in concatenation method: inference frameworks (concatenation [partitioning schemes: codon-partitioned *wpc*]), and phylogenetic signal filtering approaches (traditional metrics: evolutionary rate—EvolRate, relative composition frequency variation—nRCFV, site concordance—DVMC; novel metrics: GC-content divergence—absDiffGC and branch-length ratio—absRatioLen).

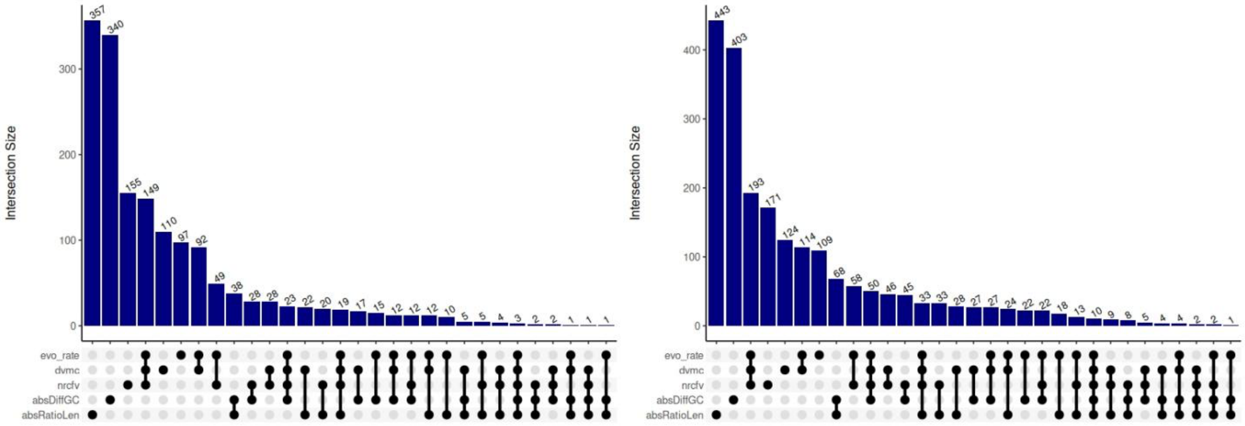


Fig. S27. The intersection among the top 500 (left panel), and 700 (right panel) most informative CDS selected from 3 traditional metrics and 2 novel metrics. The vertical bar plot displays the number of CDS shared among the five metrics. Black-filled dots connected by vertical lines indicate the intersections among the corresponding metrics. Novel filtering methods retained 76–82% unique loci in the 300-locus dataset, 68–71% in the 500-locus dataset, and 57–63% in the 700-locus dataset. Traditional methods retained 22%, 28%, and 32% unique loci (300 loci), 19%, 22%, and 31% (500 loci), and 15%, 17%, and 24% (700 loci). Although the proportion of unique loci declines with increasing dataset size, novel methods consistently retain a high level of independently selected loci.

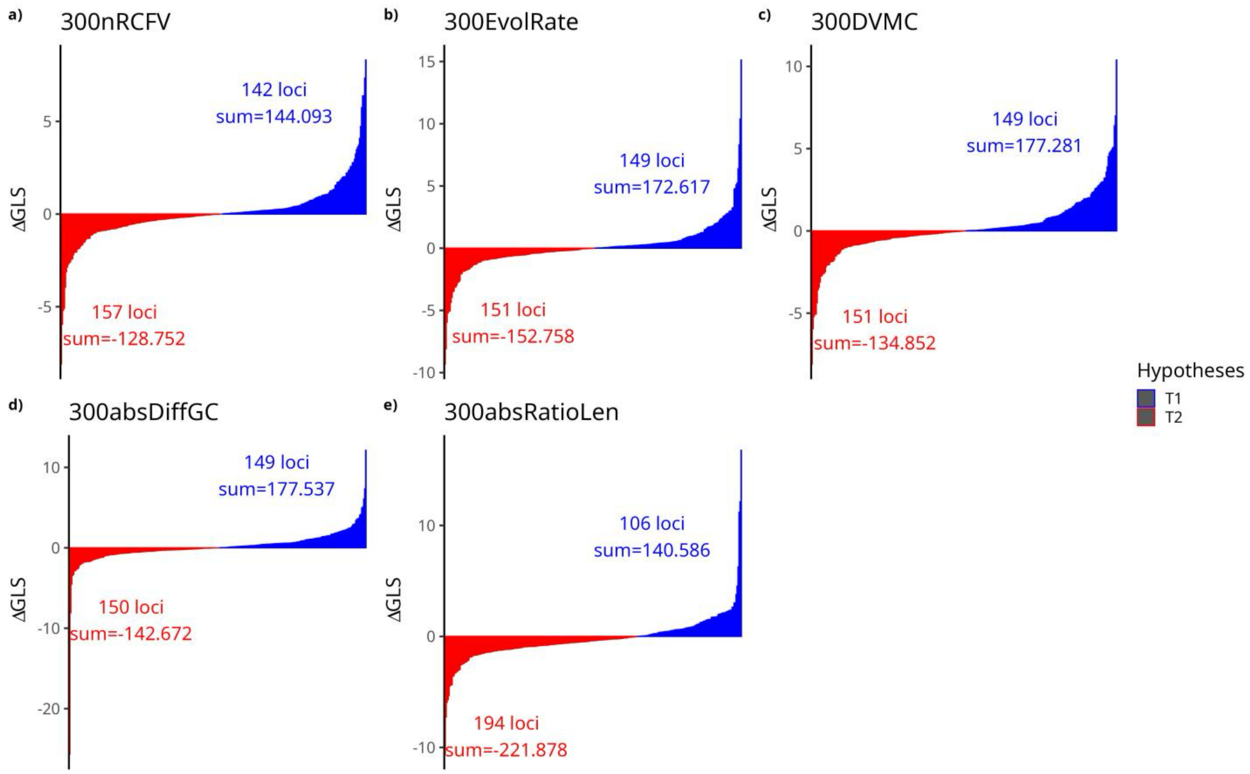


Fig. S28. Distribution of ΔGLS values across five subsets was generated based on different metric-based filtering methods. The number of loci supporting which topology and the total ΔGLS value are labeled to reflect the strength of support.

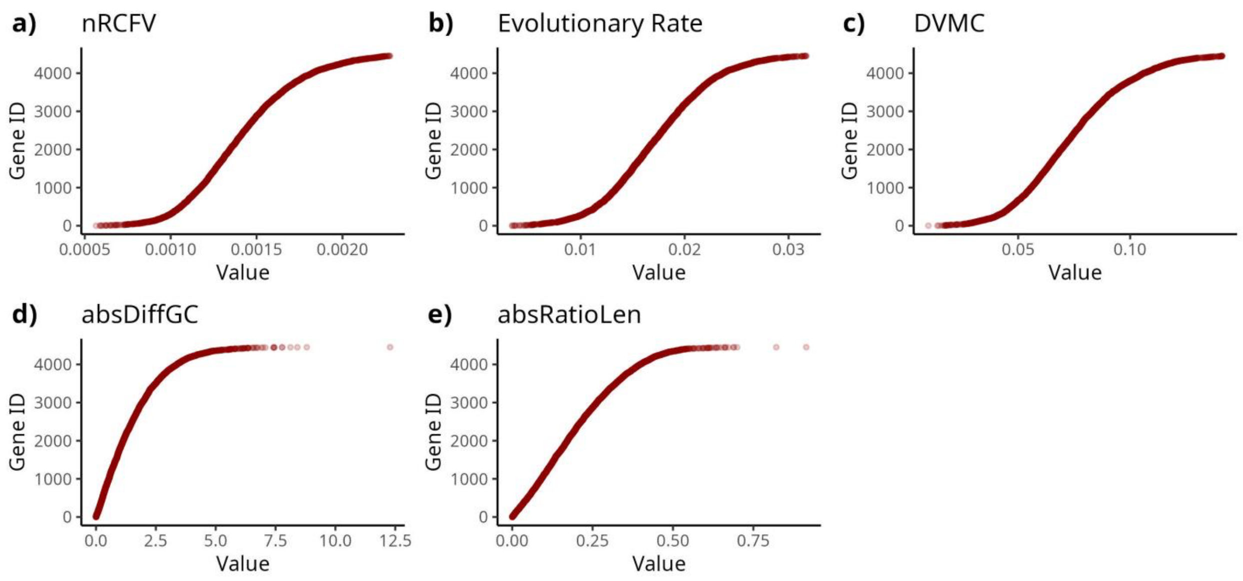


Fig. S29. Dot plots showing the distribution patterns of five metric values across the 4,452 CDS loci, sorted by value for each metric.

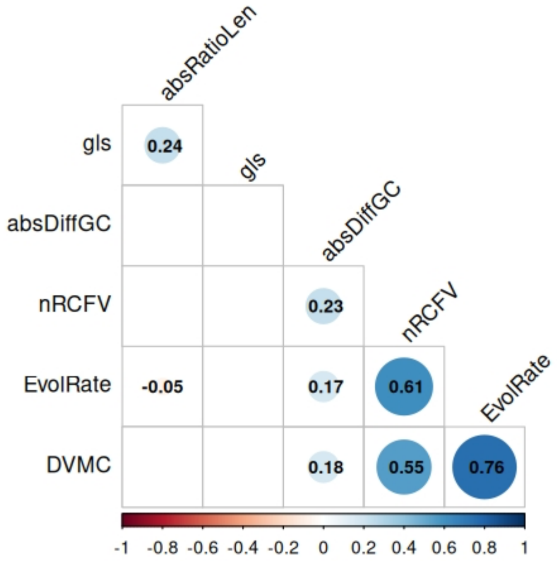


Fig. S30. Pairwise correlations among five phylogenetic signal metrics and ΔGLS.

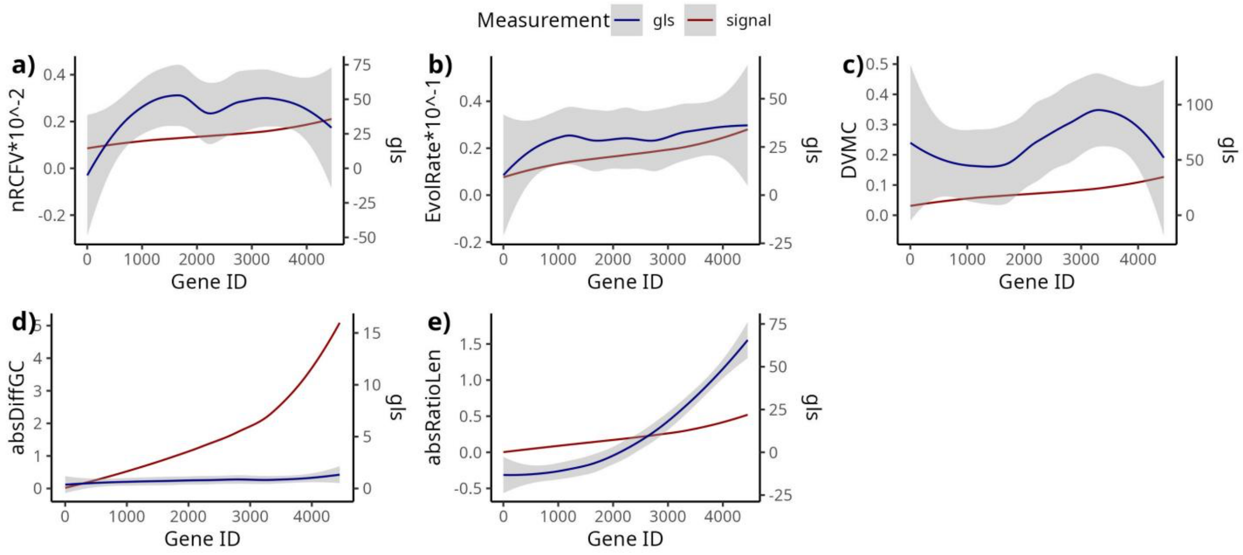


Fig. S31. Smoothed trend line showing the relationship between the number of genes (accumulated) and two y-variables—filtering metrics value (red) and ΔGLS (blue). The genes were sorted for each of the metic values from low to high (red lines). The curve was generated using the geom_smooth() function in the ggplot2 R package with 95% confidence intervals.

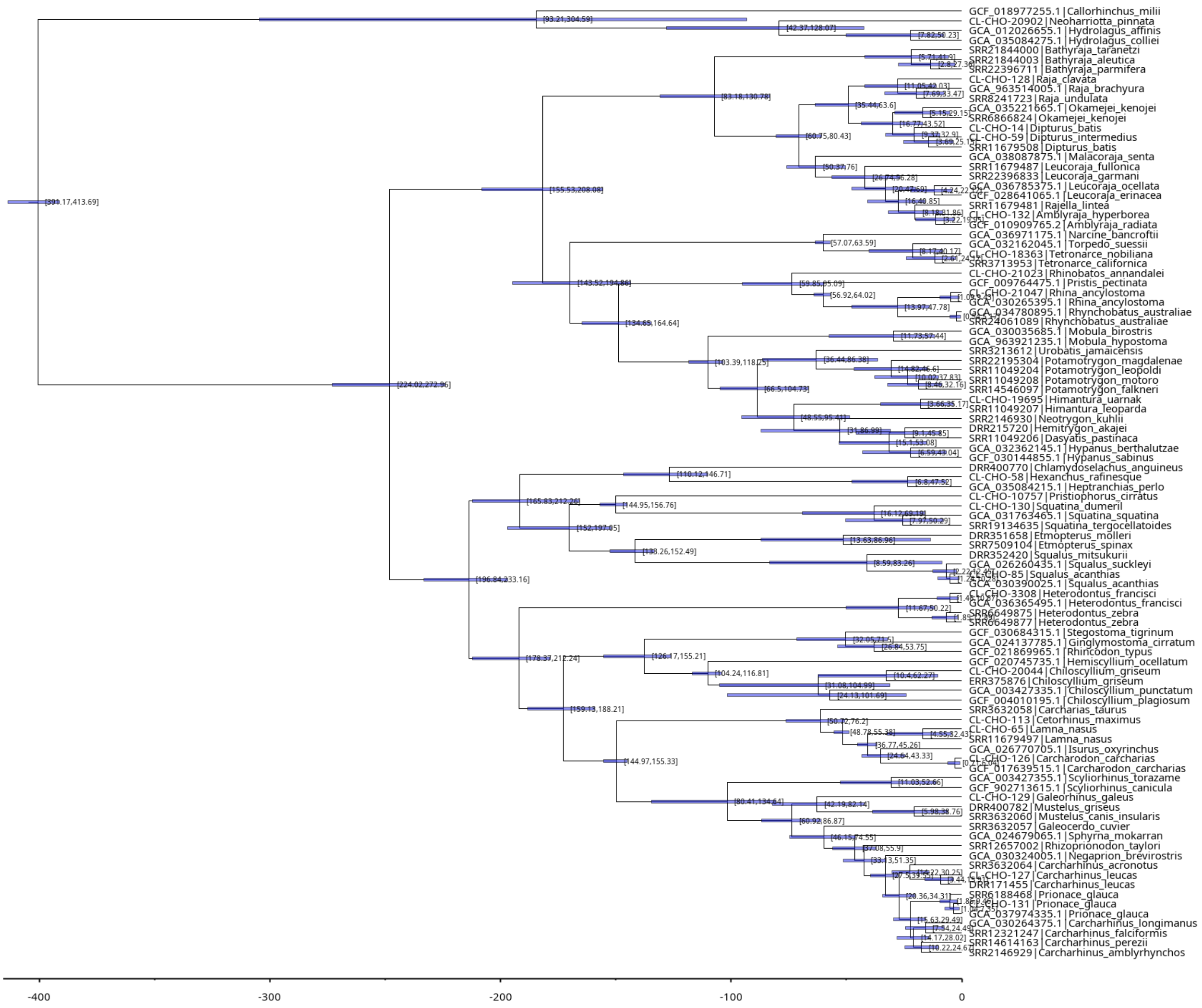


Fig. S32. Divergence time estimation for Chondrichthyes based on relaxed-clock model calibration in BEAST2. All 20 fossil calibrations are documented in Supplementary Table S8.

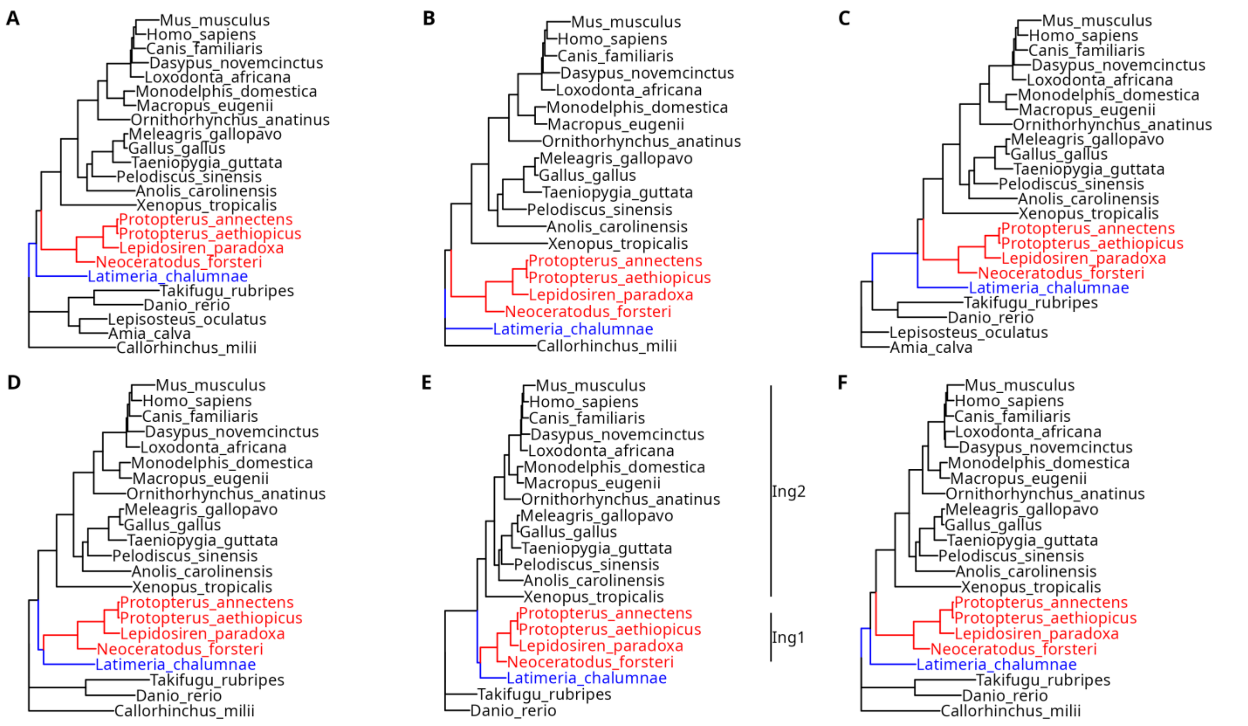


Fig. S33. Maximum likelihood phylogenies reconstructed from 4,593 amino acid alignments with no deep paralogues (Irisarri et al. 2017) under the site-homogeneous LG+G model with 1,000 ultrafast bootstrap replicates. Following the strategy of Takezaki and Nishihara (2017) to assess the effect of outgroup choice on the phylogeny of lungfish, coelacanths, and tetrapods, we retained 24 species from the original dataset and tested five outgroup configurations: (A) Chondrichthyes + ray-finned fishes (Holostei + Teleostei), (B) Chondrichthyes only, (C) ray-finned fishes only, (D) Chondrichthyes + Teleostei, and (E) Teleostei only. (F) The metric-based filtering method absRatioLen was applied to dataset E to reconstruct an additional ML tree and evaluate its impact on the inferred positions of lungfish, coelacanth, and tetrapod. In panel F, ingroup1 (lungfish) and ingroup2 (tetrapod) are labeled out. Lungfish clades are highlighted in red and the coelacanth clade in blue.
